# Improved Bayesian inference of hybrids using genome sequences

**DOI:** 10.64898/2025.12.26.696621

**Authors:** Sneha Chakraborty, Bruce Rannala

**Affiliations:** Department of Ecology and Evolutionary Biology, University of California, Los Angeles; Department of Evolution and Ecology, University of California, Davis

**Keywords:** hybrid inference, Bayesian inference, posterior predictive distribution, recombination, whole genome sequence data

## Abstract

A Bayesian hybrid inference method is developed which infers hybrids and back-crosses across two generations using sampled genomes from two populations. The method improves on that of Chakraborty and Rannala (2023) by accounting for uncertainty of population haplotype frequencies and correctly marginalizing over haplotypes while still modeling linkage and recombination across the genome. In analyses of simulated data the new method produced posterior probabilities nearly identical to the method of Chakraborty and Rannala (2023) when sample sizes were large. For small sample sizes, posterior probabilities produced by the new method tended to be lower as expected since it accounts for additional uncertainties of population haplotype frequencies. Statistical performance of the new method as measured by the ROC (Receiver Operation Characteristic) curve, appears equivalent to that of Chakraborty and Rannala (2023). The new method is applied to three recently published datasets for populations of kiwifruit (genus *Actinidia*), plateau fence lizard (*Sceloporus tristichus*) and puma (*Puma concolor*).

## INTRODUCTION

The problem of identifying recent hybrids using genomic data is fundamental to many areas of biology such as conservation genetics (Fitzpatrick *et al*., 2015), population genetics (Abbott *et al*., 2016), speciation theory (Barton, 2001), and the study of invasive species (McGaughran *et al*., 2023). Genomic datasets potentially provide powerful information for identifying hybrids but require a realistic model of hybrid transmission genetics to obtain accurate likelihoods and posterior probabilities. The Bayesian hybrid inference algorithm previously developed by Chakraborty and Rannala (2023) treats point estimates of population haplotype frequencies derived from the posterior mean of a sample as proxies for the unknown population haplotype frequencies and ignores uncertainties of these estimates in calculating likelihoods and posterior probabilities of hybrid genealogical classes. The theory developed in Chakraborty and Rannala (2023) was an improvement over an existing composite likelihood method, NewHybrids (Anderson and Thompson, 2002) in that it accounted for linkage between SNPs. We extend the method of Chakraborty and Rannala (2023) to account for the uncertainty of population haplotype frequencies by analytical integration under a conjugate prior. This produces posterior probabilities of individual assignments to hybrid genealogical classes conditional on a finite sample of individuals from each source population. Chakraborty and Rannala (2023) also used a method to marginalize over haplotypes (when 2 or more recombinations occur) that was biologically unrealistic. Essentially, they treated each contiguous sub-haplotype as if it were independently sampled from the population, which is not the case; the sub-haplotype (including all segments whether contiguous or not) should be sampled jointly. We also present here a new more realistic algorithm for correctly marginalizing over haplotypes. We use simulations to explore the effects of population sample sizes on assignment probabilities and compare the posterior assignments with those obtained using the Chakraborty and Rannala (2023) method. We also use the new method to analyze example empirical datasets for kiwifruit, lizards and puma.

## THEORY

We consider a scenario where a sampled individual may be admixed between two populations A and B. Samples of pure individuals are available from populations A and B which we refer to as the *reference samples*. Here we develop the theory for a single chromosome; this is extended later by adding an additional subscript to indicate the particular chromosome being considered. Let **f** ^*k*^ = {*f*^*k*^(*h*)} be the haplotype frequencies in population *k* ∈ {*A, B*}, where *f*^*k*^(*h*) is the frequency of the *h*-th distinct haplotype. Let *N*_*k*_ be the number of diploid individuals in a phased reference sample from population *k*. It is assumed that *H* distinct haplotypes exist, each occurring in both populations. The set of distinct haplotypes compatible with genotypes observed in all sampled individuals (from both populations) provides an estimate of *H*. With no prior information, we assign equal prior probability density to the haplotype frequencies in each population (A or B), so the frequencies are **f** ^*k*^ ∼ Dirichlet(1/*H*). The prior probability density of haplotype frequencies (i.e., before collecting reference samples) in population *k* is

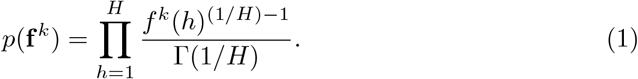

Let **n** = {**n**_*A*_, **n**_*B*_} where **n**_*k*_ = {*n*_1*k*_, …, *n*_*Hk*_} and *n*_*hk*_ is the observed number of copies of the *h*th distinct haplotype in a reference sample from population *k*. The probability mass of **n**_*k*_ conditioned on the haplotype frequencies follows a Multinomial distribution given by

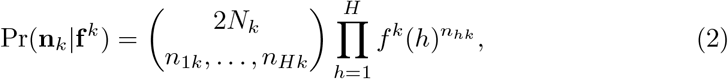

where 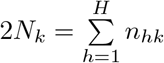 is the total number of haplotypes sampled in population *k*. The posterior density of the haplotype frequencies, conditioned on the haplotypes observed in a reference sample from population *k*, follows a Dirichlet distribution, since the Dirichlet is a conjugate prior to a Multinomial distribution. The posterior probability density of haplotype frequencies is

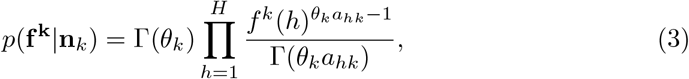

where, *θ*_*k*_ = 1 + 2*N*_*k*_ and 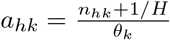. Note that 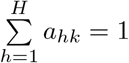.

### Posterior Predictive Distribution

To calculate the likelihoods of the individual diplotypes, accounting for uncertainties of haplotype frequencies, we require the *posterior predictive distribution*. The posterior predictive distribution of the observed haplotype counts, **ñ** = {*ñ*_1_, …, *ñ*_*H*_} from a sampled individual diplotype, conditioned on the reference sample haplotype counts **n**_*k*_ from population *k* is

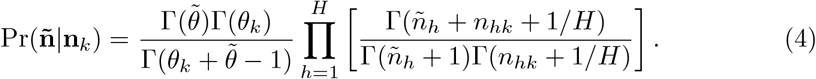

The log probability is

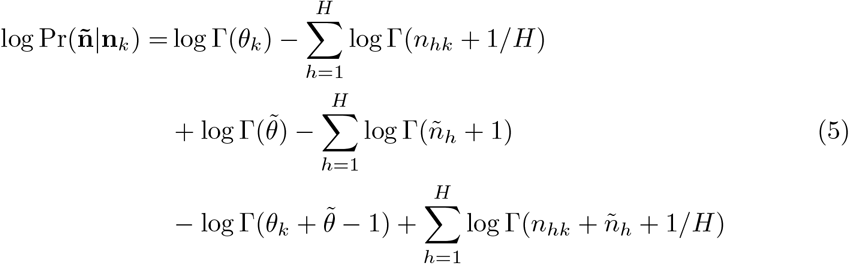

where, 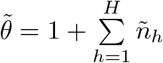

### Data and Parameters

Following Chakraborty and Rannala (2023), we consider a diploid individual with *K* chromosomes. Chromosome *i* contains *L*_*i*_ loci with phased biallelic single-nucleotide polymorphisms. The maternally (M) and paternally (P) inherited chromosomes are defined in the form of matrices:

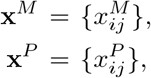

where 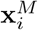 and 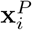 are haplotypes for maternally and paternally inherited copies of chromosome *i* and 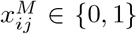 (or 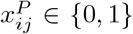) is the allele (coded as 0,1) present at the *j*th SNP locus on the maternally (*M*) (or, paternally (*P*)) inherited copy of chromosome *i*. The diplotype is the pair of haplotypes on homologous chromosomes, denoted as **x** = {**x**^*M*^, **x**^*P*^}.

For chromosome *i*, we assume that *H* distinct haplotypes exist, each occurring in both populations A and B. The reference population haplotypes are defined in terms of *H* × *L*_*i*_ matrices:

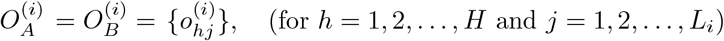

where 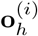 is the *h*-th haplotype of chromosome *i* and 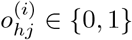 is the allele (coded as 0,1) present at the *j*-th SNP locus on the chromosome *i*.

### Likelihoods for Genealogical Classes

Two populations hybridizing over two generations will allow an individual to be classified into one of 6 distinct genealogical classes based on the number and arrangement of founder population originations. The 4 founders in this framework are considered to have pure ancestry from a reference population and the individual to be classified is at the base of the pedigree (see Figure 1 of Chakraborty and Rannala, 2023). The 6 genealogical classes are as follows: **a** and **d** are purebreds; **b** and **e** are backcrosses; **c** is an F1 hybrid and **f** is an F2 hybrid. We use the term genealogical class and model interchangeably. Here, we present formulas for calculating the likelihood of an individual’s diplotype, **x** = {**x**^*M*^, **x**^*P*^} under each of the 6 possible genealogical classes. Let *G* = {*a, b, c, d, e, f*} denote the set of all genealogical classes where *g* ∈ *G*. We define the indicator function for the *i*-th chromosome,

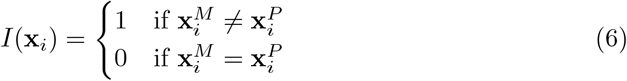

where equality denotes identical haplotypes and inequality differences between haplotypes at one or more positions. We define a bijective function ***ϕ***(**x**_*i*_) : **x**_*i*_ ↦ **ñ** such that ***ϕ***(**x**_*i*_) = [*ϕ*_1_(**x**_*i*_), …, *ϕ*_*H*_ (**x**_*i*_)] where,

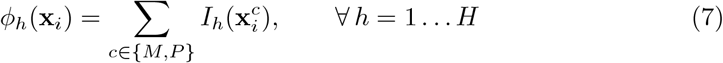

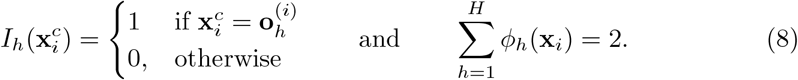

**Fig. 1.**
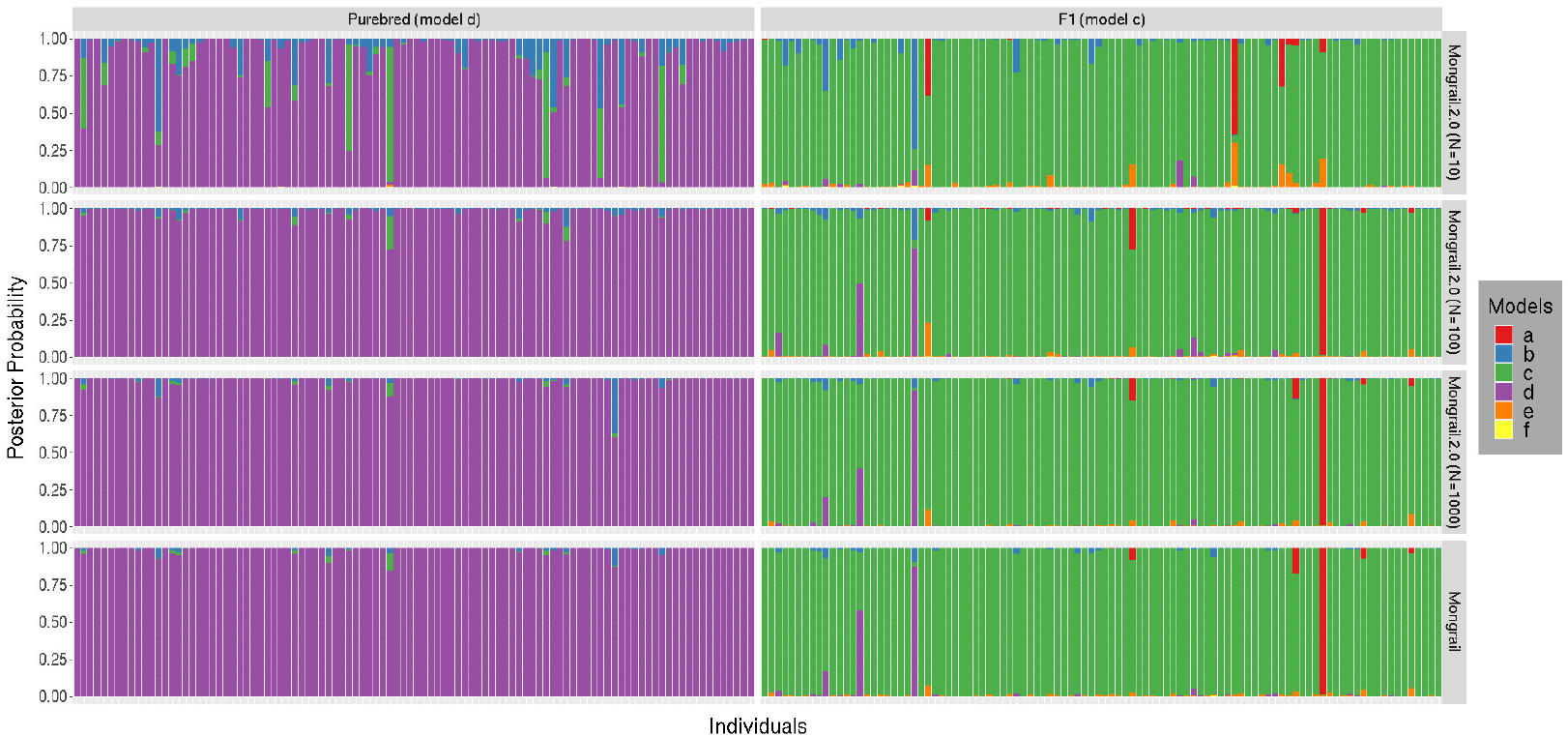
Distribution of posterior probabilities for 100 individuals simulated under genealogical classes **d** (left column) and **c** (right column) using the following set of simulation parameters: *K* = 20, *L* = 10, *R* = 50, *h* = 5, *c* = 0.1, *α* = 1. Posterior probabilities using Mongrail with known population haplotype frequencies is shown in bottom plot (4th row) and Mongrail 2.0 for different values of Multinomial sample counts *N* = 10, 100, 1000 in the first three rows respectively. The six genealogical classes are as follows: **a**-pure population B, **b**-backcross with population A, **c**-F1 hybrid, **d**-pure population A, **e**-backcross with population B, **f** -F2 hybrid.

Similarly, we define a unit vector function 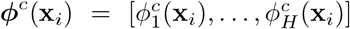, where 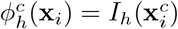. Below we define the log-probability of the likelihood of the diplotype for any chromosome under each of the 6 genealogical classes. Note that the functions *U* and *Q*, which appear below and determine haplotype probabilities involving recombination, are quite complex and are therefore defined in separate appendices (Appendix A and Appendix B respectively). The definition of *Q*(**z** | *d*_*i*_, *r*) is the same as that of (Chakraborty and Rannala, 2023) but for ease of reference is restated in Appendix B.

#### Model (a): Purebred population B

Both chromosomes (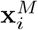 and 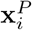) come from Population B. The log-likelihood of the *i-*th diplotype for an individual is equivalent to the log-probability of the predicted counts of the two haplotypes as given by

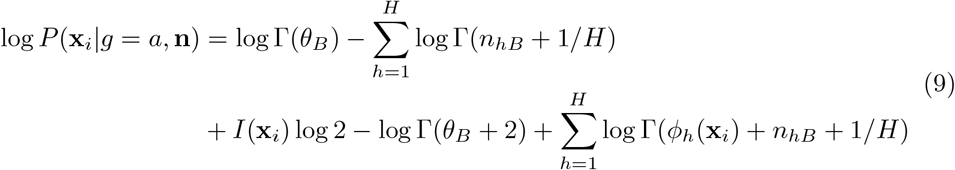

#### Model (b): Backcross with Population A

One chromosome comes from Population A and the other chromosome is a recombinant.

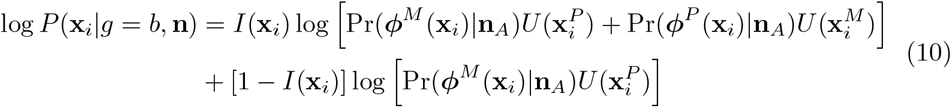

where, 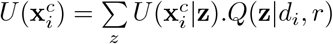, and

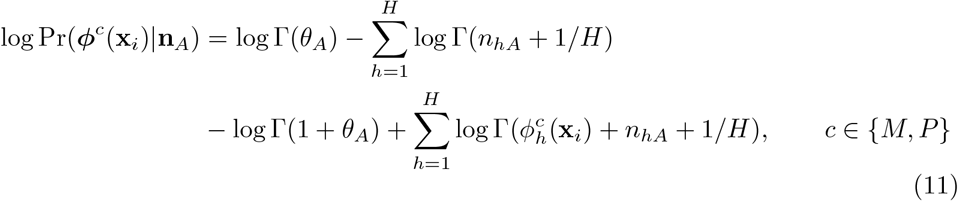

#### Model (c): F1 hybrid

One chromosome comes from Population A and the other from Population B

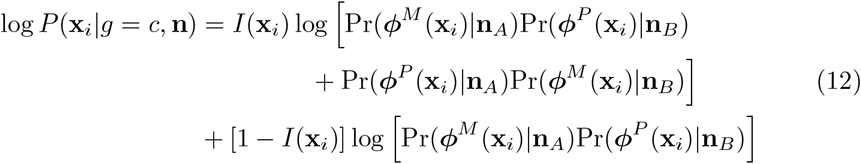

where,

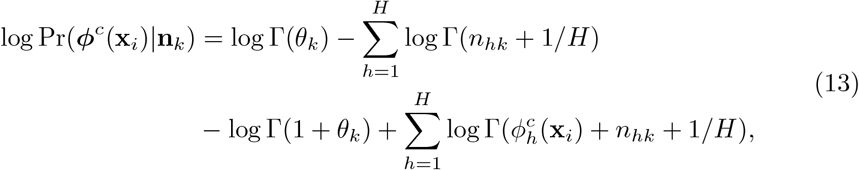

*c* ∈ {*M,P*}, *k* ∈ {*A,B*}

#### Model (d): Purebred population A

Both chromosomes (*x*^*M*^ and *x*^*P*^) come from Population A. The log-probability of the predicted counts of the two haplotypes is given by

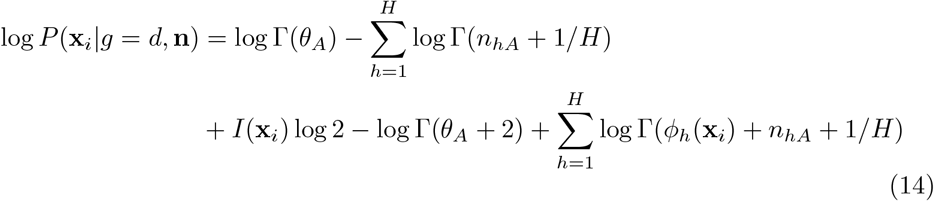

#### Model (e): Backcross with Population B

One chromosome comes from Population B and the other chromosome is a recombinant.

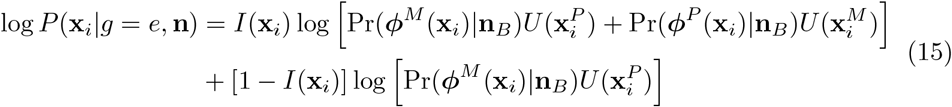

where, 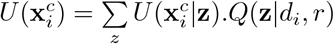, and

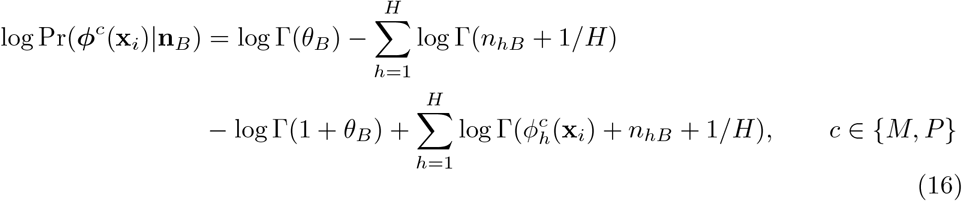

#### Model (f): F2 hybrid

Both chromosomes are recombinants.

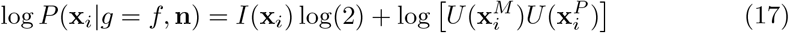

where, 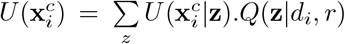, *c* ∈ {*M, P* }. Therefore the likelihood of a diplotype, **x** = {**x**^**M**^, **x**^**P**^**}** for an individual with *K* chromosomes under the *g*-th genealogical class is given by,

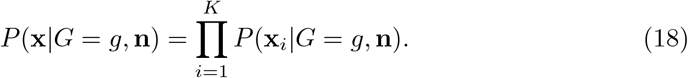

### Posterior Probability of Genealogical Classes

The posterior probability that an individual with diplotype **x** belongs to the *g*-th genealogical class is

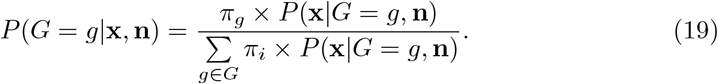

where, an individual belongs to genealogical class *g* with prior probability *π*_*g*_ with *g* ∈ *G*

## SIMULATION STUDY

We used simulated data to compare the Mongrail program (Chakraborty and Rannala, 2023) with our new method implemented as Mongrail 2.0 which relaxes the assumption of known population haplotype frequencies and correctly calculates marginal haplotype probabilities under recombination. We use a subset of the simulated datasets from Chakraborty and Rannala (2023), fixing the following combination of parameters: *K* = 20, *L* = 10, *c* = 0.1, *α* = 1. We examined the relative performance of the two methods under different values of *R* (1cM or 50cM) and *h* (5 or 15). See Supplementary Article section S1.1 for details. Chakraborty and Rannala (2023) used a Dirichlet distribution to generate a diverse set of haplotype frequency distributions which were reused in the current study. Diplotypes for hybrid individuals were simulated based on the simulated haplotypes and their corresponding frequencies. For Mongrail 2.0, we use these previously generated frequencies as parameters for a Multinomial distribution to generate samples of haplotypes from the reference populations. We considered multinomial sample sizes of *N* = 10, 100, 1000 to generate different reference population datasets. For each simulated individual we either used Mongrail posterior probabilities computed in our previous paper (using population frequencies) or computed posterior probabilities using the posterior mean from the sample counts. We applied our new method to the same set of simulated individuals using the simulated reference samples to compute the posterior probabilities. We use the following three metrics to study the statistical performance of Mongrail and Mongrail 2.0:

1. Influence of sample size on posterior probabilities
2. AUC-ROC curve analysis

### Influence of sample size on posterior probabilities

Here we compare the posterior probabilities previously computed using Mongrail and known population haplotype frequencies (Chakraborty and Rannala, 2023) with those of Mongrail 2.0 using different multinomial sample sizes (*N* = 10, 100, 1000). Only the stacked bar plots for 100 individuals from each of the six genealogical classes are shown as other results are essentially similar. Figure 1 shows that, irrespective of population sample size (*N*), both methods assign higher posterior probabilities to the true genealogical class of an individual for purebreds and F1 hybrids. The posterior probabilities are lower and distributed across more genealogical classes for Mongrail 2.0 when *N* = 10 and *N* = 100, reflecting the additional uncertainty when haplotypes sampling from the reference populations is properly accounted for. This effect is particularly notable for backcrosses and F2 hybrids.

Figure 2 shows the results for backcross and F2 hybrid individuals as the sample size increases from *N* = 10 to 1000 (moving top to bottom, first three rows). The magnitude of posterior probabilities produced by Mongrail 2.0 for the correct genealogical class (backcrosses and F2 hybrids) increases. The performance of Mongrail 2.0 under a multinomial sample size of *N* = 1000 (third row) is very similar to Mongrail used with known frequencies (last row). For a small sample size of *N* = 10 (first row), Mongrail 2.0 assigns more posterior probability to alternative genealogical classes as expected. It is evident that as the model complexity increases (from figure 1 to figure 2), the difference between the two methods Mongrail and Mongrail 2.0 is more pronounced. However, with a sample size of *N* = 1000, the haplotype count data simulated for use with Mongrail 2.0 is close to the expected counts under the true haplotype frequency distribution as used by Mongrail. Thus the methods are comparable for *N* = 1000.

**Fig. 2.**
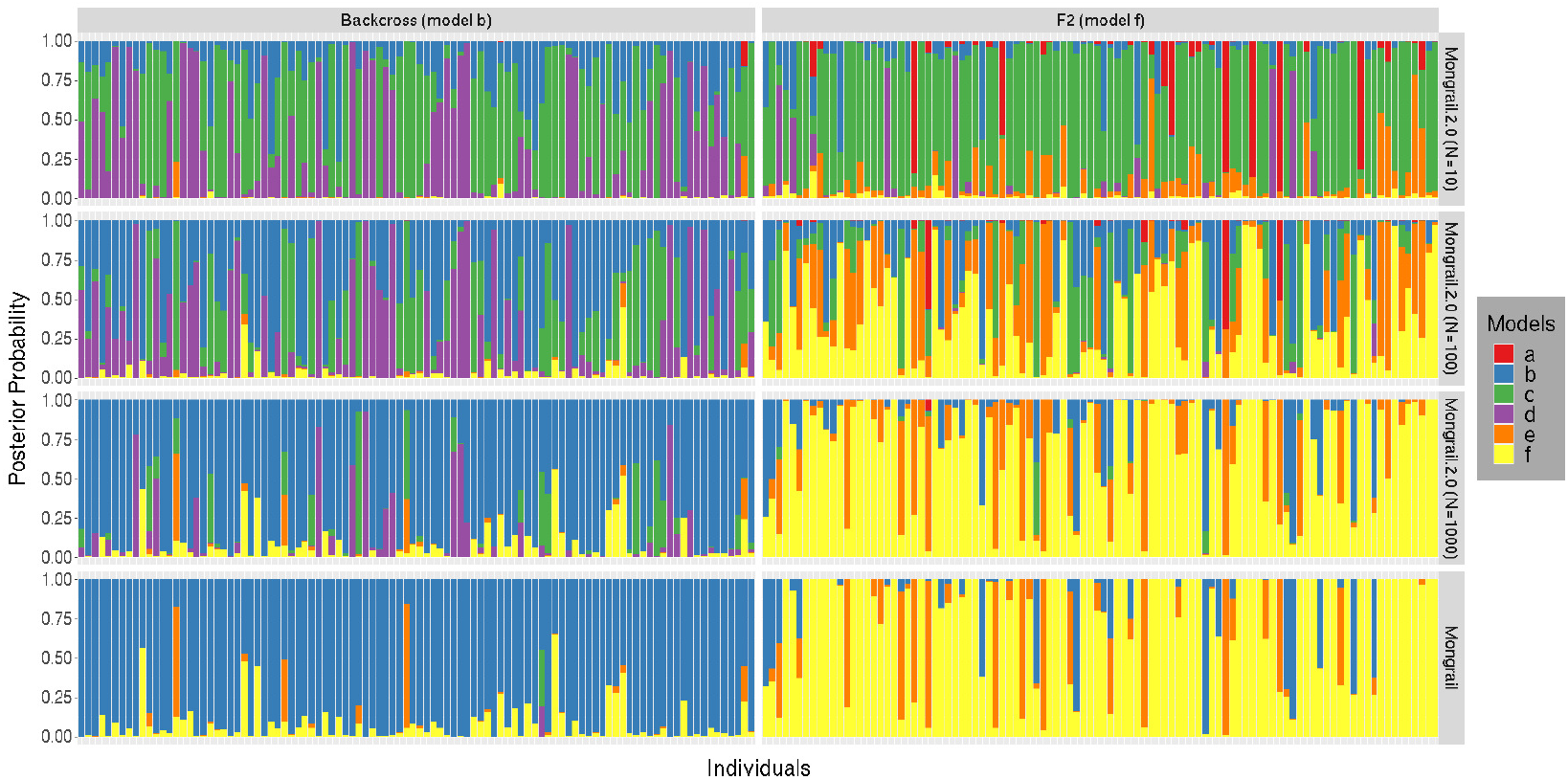
Distribution of posterior probabilities for 100 individuals simulated under genealogical classes **b** (left column) and **f** (right column) using the following set of simulation parameters: *K* = 20, *L* = 10, *R* = 50, *h* = 5, *c* = 0.1, *α* = 1. Posterior probabilities using Mongrail with known population haplotype frequencies is shown in bottom plot (4th row) and Mongrail 2.0 for different values of Multinomial sample counts *N* = 10, 100, 1000 in the first three rows respectively. The six genealogical classes are as follows: **a**-pure population B, **b**-backcross with population A, **c**-F1 hybrid, **d**-pure population A, **e**-backcross with population B, **f** -F2 hybrid.

### AUC-ROC curve analysis

Here we examine the performance of the two methods to classify the different genealogical classes for different classification thresholds using the AUC-ROC curve analysis. The greater the area under the ROC curve (AUC) the better the performance. We compare the AUC values of Mongrail (posterior probabilities computed using the posterior mean from the sample counts) with Mongrail 2.0 under different multinomial sample sizes (*N* = 10, 100 or 1000), recombination frequency (*R* = 1cM or 50cM) and distinct haplotype counts per chromosome for each population (*h* = 5 or 15). We are interested in the interaction of the different parameters (*N, R* and *h*) and their effect on classification power of the two methods. We plot the AUC values against increasing values of *N* as multi-line plots, where line colors represent different methods (Mongrail 2.0 and Mongrail) and line types (solid or dashed) different recombination frequencies (*R* = 1cM or 50cM). The results are shown in figure 3 where the top and bottom panels represent the cases for *h* = 15 and *h* = 5, respectively.

**Fig. 3.**
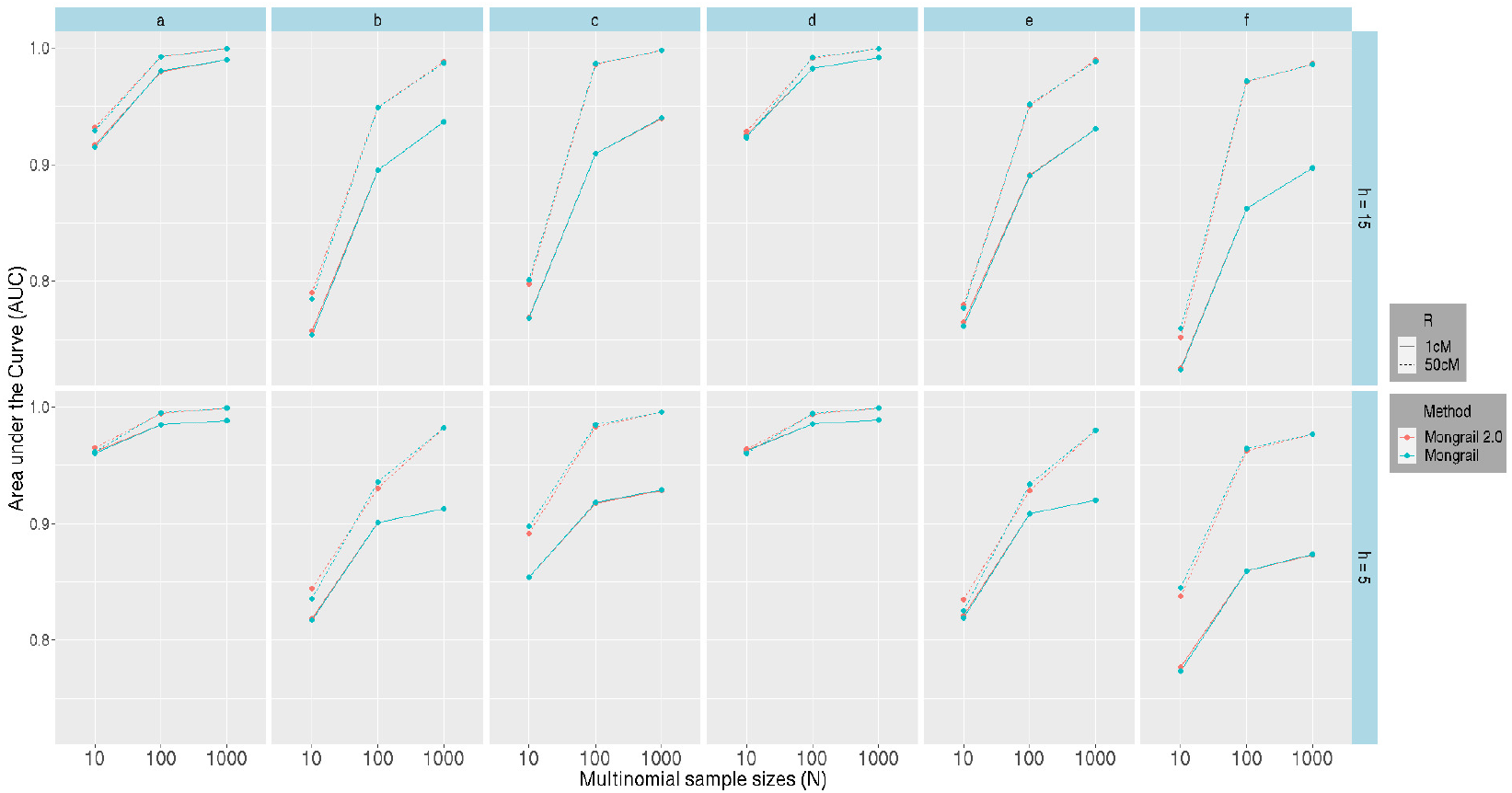
AUC (Area under the curve) values (y-axis) plotted against different multinomial sample sizes *N* (x-axis) for Mongrail 2.0 (red line) and Mongrail (blue line). The plot is based on 10,000 individuals simulated using parameters: *K* = 20, *L* = 10, *c* = 0.1, *α* = 1 and either expected recombination frequency of *R* = 1cM (solid linetype) or *R* = 50cM (dotted linetype). The first and second row corresponds to number of distinct haplotypes per chromosome with values *h* = 15 and 5 respectively. Results for the six genealogical classes (**a**-**f**) are shown from left to right in both rows. The 6 genealogical classes are as follows: **a**-pure population B, **b**-backcross with population A, **c**-F1 hybrid, **d**-pure population A, **e**-backcross with population B, **f** -F2 hybrid.

Figure 3 shows that irrespective of the method and any values of *R* or *h*, the AUC increases as the sample size increases from *N* = 10 to 1000. The performance of the two methods improves as recombination frequency increases from *R* = 1 cM to 50 cM (solid to dashed line) as expected. Performance of the alternate methods is almost indistinguishable across all six genealogical classes, especially for sample sizes *N* = 100 and 1000, regardless of values for *R* or *h*. We also observe that for *N* = 10 and *N* = 100, AUC values are higher for *h* = 15 relative to *h* = 5 across all genealogical classes. The difference is negligible for *N* = 1000 and more pronounced for *N* = 10. This suggests that with a small sample size there is limited information about population haplotype frequencies (and predicted haplotype counts) when the number of distinct haplotypes increases. This uncertainty affects both methods; Mongrail 2.0 is influenced through the distribution of predicted haplotype counts) and Mongrail is affected because the posterior mean is based on the sample counts. The AUC values in Figure 3 summarize the combined impact of the parameters *h, R* and *N* on the power of the two methods to detect genealogical classes. For further details on individual parameter effects see the individual ROC curves (Figures S1-S6) in the Supplementary Article section S1.2.

## EMPIRICAL ANALYSIS

To assess the performance of the new method and the Mongrail 2.0 program when applied to empirical datasets for a range of organisms we analyzed three published hybridization datasets: kiwifruit (Yu *et al*., 2023), plateau fence lizards (Leaché *et al*., 2025) and Florida pathers (Aguilar-Gómez *et al*., 2025).

### Kiwifruit

We analyzed a dataset consisting of two deeply divergent kiwifruit species (*Actinidia eriantha* and *Actinidia hemsleyana*) and their hybrids using Mongrail 2.0. Actinidia, a dioecious variety of plant is mainly found in East and South Asia. *A. eriantha* is quite widespread unlike *A. hemsleyana* which occurs mainly in Eastern China. *Actinidia zhejiangensis* is thought to be a homoploid hybrid between *Actinidia eriantha* and *Actinidia hemsleyana*. This species is quite rare and is found in Jiangxi, Zhejiang and Fujian provinces in China. Yu *et al*. 2023) specifically chose these species to study the evolutionary consequence of homoploid speciation.

The dataset consists of 51 individuals 21 *A. hemsleyana*, 19 *A. eriantha* and 11 *A. zhejiangensis*). Whole genome resequencing data were assembled against *A. eriantha* genome and variant calling were performed to generate a Variant Call Format (VCF) data file. We used BCFtools to filter out missing sites and obtained a dataset with 5,345,755 SNPs on 29 chromosomes. We used Beagle 5.1 to phase the data for the two parental populations (*Actinidia eriantha* and *Actinidia hemsleyana*). See Supplementary Article section S2.1 for more details on recombination rate and markers. We used Mongrail 2.0 to calculate the posterior probabilities of genealogical classes for these data. In this framework *A. eriantha* is treated as population A and *A. hemsleyana* as population B. The genealogical classes are therefore as follows:

- Model a - *A. hemsleyana*
- Model b - Backcross with *A. eriantha*
- Model c - F1 hybrid (presumptive *A. zhejiangensis*)
- Model d - *A. eriantha*
- Model e - Backcross with *A. hemsleyana*
- Model f - F2 hybrid

Figure 4 shows the posterior probability distribution of 11 presumed *A. zhejiangensis* individuals. Mongrail 2.0 produces very high posterior probability for 10 out of 11 individuals. Based on a classification threshold of 0.95, 10 individuals were assigned to specific genealogical classes with posterior probability higher than 0.95. Seven were identified as F1 hybrids (Individuals: JNZJ-1, JNZJ-3, LQZJ-1, TSZJ-1, WCZJ-1, WCZJ-2, YP-1), two as F2 hybrids (Individuals: ASX-1, JNZJ-5) and one as back-crossed with *A. eriantha* (Individual: LSZJ-1). For direct comparison, we also applied Mongrail to this dataset (see Supplementary Section S3, Figures S7). Results were consistent across methods for all individuals.

**Fig. 4.**
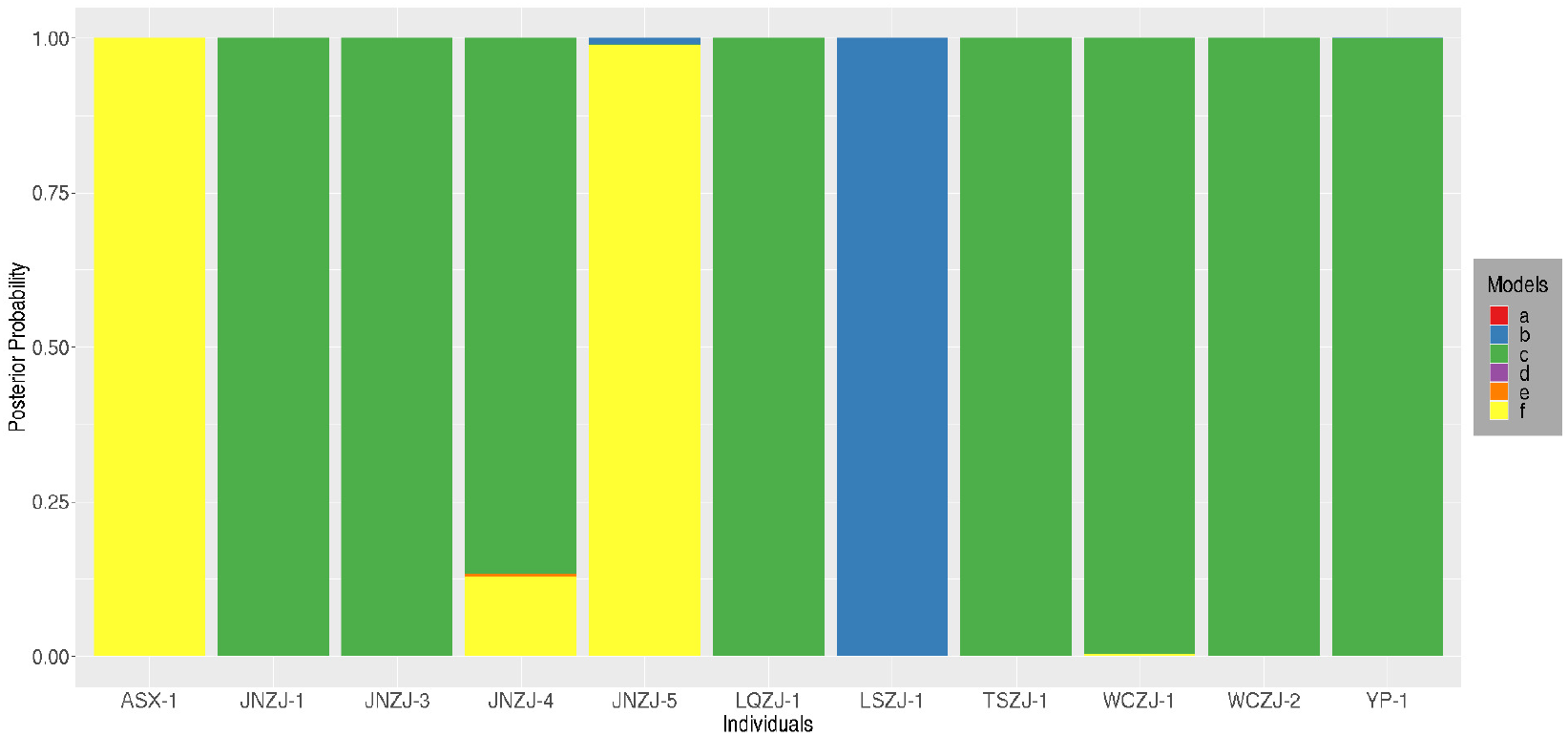
Stacked bar plot showing the distribution of posterior probabilities for 11 presumed *A. zhejiangensis* individuals. The genealogical classes are : **a** - *A. hemsleyana*, **b** - Backcross with *A. eriantha*, **c** - F1 hybrid, **d** - *A. eriantha*, **e** - Backcross with *A. hemsleyana* and **f** - F2 hybrid.

### Lizard

Next we analyzed plateau fence lizards (*Sceloporus tristichus*) sampled along a transect between Great Basin Grassland (north) and Great Basin Conifer Woodland (south) in eastern Arizona (USA) (collected in 2022 by Leaché *et al*. (2025)). The lizards found in grasslands (north) and juniper woodlands (south) are the two distinct parental populations. A hybrid zone is located in an ecotone where the vegetation transitions from grassland to juniper habitat between the parental populations. The sampling transect includes eight sites (Holbrook, Fivemile Wash, Washboard Wash, Woodruff, Canoncito, Sevenmile Draw, Snowflake and Show Low) spanning a distance of 63.5 km where the hybrid zone is located primarily between the cities of Holbrook (north) and Show Low (south). We refer to parental grassland and juniper lizard populations as Holbrook and Show Low populations respectively. The remaining samples from six different sites are treated as potential hybrids. Leaché *et al*. (2025) collected these data to study the temporal introgression dynamics of the two populations sampled along the linear transect.

The dataset consists of 78 individuals (7 Holbrook, 11 Show Low and 60 transect samples). Among the 60 individuals sampled from the transect, 11 were from Fivemile Wash, 10 from Washboard Wash, 9 from Woodruff, 10 from Canoncito, 10 from Sevenmile Draw and 10 from Snowflake. Leaché *et al*. (2025) collected SNP data using double-digestion restriction site-associated DNA sequencing (ddRADseq) protocol, aligning the reads against a reference genome of *Sceloporus tristichus* (Bedoya and Leaché, (2021) and generated a Variant Call Format (VCF) data file. We used BCFtools (version 1.19-43) to filter out missing data, retaining only biallelic sites leaving 714 SNPs on 11 chromosomes. We used Beagle 5.1 to phase the data for the two parental populations (Holbrook and Show Low). See Supplementary Article section S2.2 for more details on recombination rate and markers. Mongrail 2.0 was used to calculate posterior probabilities of genealogical classes for individuals. In this framework Holbrook is treated as population A and Show Low as population B. The genealogical classes are therefore as follows:

- Model a - Show Low
- Model b - Backcross with Holbrook
- Model c - F1 hybrid
- Model d - Holbrook
- Model e - Backcross with Show Low
- Model f - F2 hybrid

Figure 5 shows the posterior probabilities of genealogical classes for each individual from the transect region. Most individual posterior probabilities are distributed primarily over three hybrid classes (F2 and backcrosses). Genealogical class **b** which is a backcross with Holbrook (color blue in Figure 5) has increased posterior probabilities in many individuals at the sites sampled from Fivemile Wash, Washboard Wash, Woodruff and Canoncito. By contrast, this pattern changes drastically for lizards sampled from Sevenmile Draw and Snowflake where the color red (genealogical class **a**: pure Show Low) and orange (genealogical class **e**: backcross with Show Low) is more dominant. Genealogical class assignment remains uncertain for most individuals. Only 6 out of 60 transect sampled individuals has a posterior probability higher than 0.95 for a specific genealogical class. Classification decisions based on a posterior probability threshold of 0.95 are shown in table 1. Three of 6 individuals are classified as F2 hybrids (Individuals: Five 5201, Wash 5192, Wood 5227), one as pure Holbrook (Individual: Wash 5253) and two as pure Show Low (Individuals: Nsno 5235, Snow 5264). The remaining 51 transect sampled individuals cannot be assigned with certainty to any of the genealogical classes. To allow direct comparison, we also applied Mon-grail to this dataset (see Supplementary Section S3, Figures S8-S13). Results were qualitatively similar for both methods, except for one individual (Nsno 5235, Figure S12).

**Table 1.** Assignment of 6 transect sampled plateau fence lizards.

| Individual Index | Sampling Location | Posterior Probability | Genetic Identification | Chosen Model |
| --- | --- | --- | --- | --- |
| Five_5201 | Fivemile Wash | 0.986 | F2 | f |
| Wash_5192 | Washboard Wash | 0.95 | F2 | f |
| Wash_5253 | Washboard Wash | 0.95 | Pure Holbrook | d |
| Wood_5227 | Woodruff | 0.989 | F2 | f |
| Nsno_5235 | Sevenmile Draw | 0.979 | Pure Show Low | a |
| Snow_5264 | Snowflake | 0.984 | Pure Show Low | a |

**Fig. 5.**
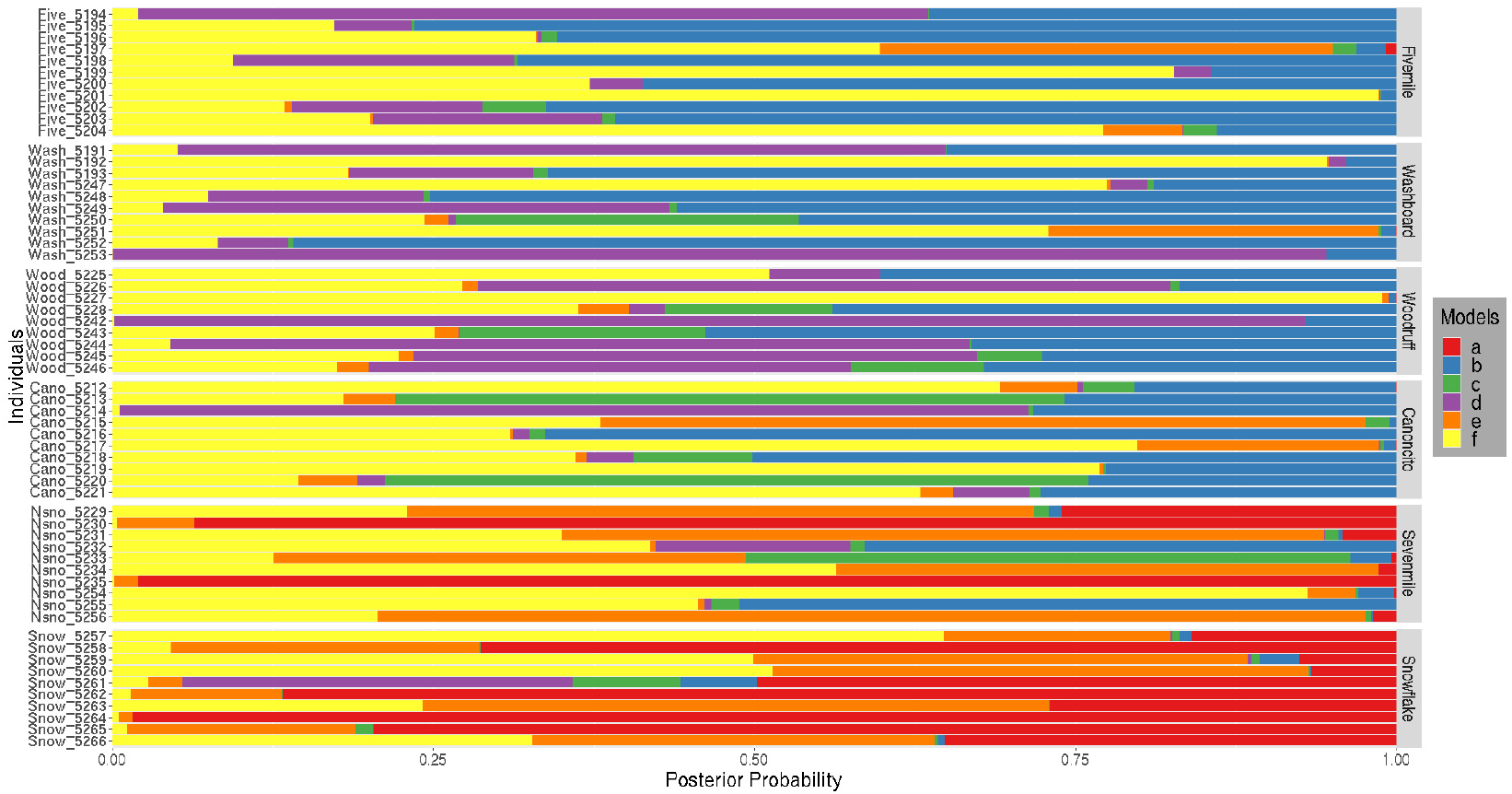
Horizontal stacked bar plot showing the distribution of posterior probabilities for 60 plateau fence lizards (*Sceloporus tristichus*) sampled from the transect. Individuals are arranged according to the sampled sites along the transect. Panels correspond to the six sites - Fivemile Wash, Washboard Wash, Woodruff, Canoncito, Sevenmile Draw and Snowflake which are ordered latitudinally. The genealogical classes are : **a**-pure Show Low, **b**-backcross with Holbrook, **c**-F1 hybrid, **d**-pure Holbrook, **e**-backcross with Show Low, **f** -F2 hybrid.

### Puma

Lastly we analyze a dataset comprising pumas (*Puma concolor*) found in southern Florida, also called Florida panthers. Florida panthers were once common in the US southeast but urban expansion and bounty hunting decimated the population, reducing their range to hardwood swamps and cypress prairies of south and central Florida. The species was listed as “endangered” in 1967 by the U.S. Fish and Wildlife Service under the Endangered Species Act. By the 1990s this small and isolated population of Florida panthers were suffering from reproductive failure and phenotypic defects due to inbreeding and less than 30 individuals remained in the wild. To rescue this population eight female panthers from Texas were released in southern Florida in 1995, five of which were successful in producing offspring. This introduction led to a four-fold increase in population size. Currently there are between 120 and 230 adult and subadult Florida panthers in the wild.

This conservation strategy for increasing the genetic diversity and fitness of a shrinking population is termed genetic rescue. Aguilar-Gómez *et al*. (2025) sequenced 29 post-rescue Florida panthers to study the long-term genomic consequences of genetic rescue. They also included previously published whole genome sequence data from two parental populations (Texas and pre-rescue Florida panthers) and two known F1 hybrids. The Texas population was represented by genome sequences collected from the five female Texas panthers originally introduced in 1995 (Ochoa *et al*., (2019). The pre-rescue Florida panther population was represented by genome sequences of 2 individuals sampled by Ochoa *et al*. (2019) and 2 sampled by Saremi *et al*. (2019). The two known F1 hybrids were from Ochoa *et al*. (2019). We applied Mongrail 2.0 to analyze this dataset consisting of 5 Texas panthers, 4 pre-rescue Florida panthers and 31 post-rescue Florida panthers. Aguilar-Gómez *et al*. 2025) aligned these genomes against a reference genome for *Puma yagouaroundi* (an outgroup) and performed variant calling to generate a Variant Call Format (VCF) file. We filtered out missing data using BCFtools (version 1.19-43) and selected scaffolds larger than 20 Mb to produce a dataset with 1,107,855 biallelic SNPs across 34 scaffolds. We used Beagle 5.1 to phase the data for the two parental populations (Texas and pre-rescue Florida panthers). See Supplementary Article section S2.3 for more details on recombination rate and markers. Mongrail 2.0 was used to calculate posterior probabilities of the genealogical class for each post-rescue individual. In this framework, pumas from Texas are treated as population A and pre-rescue Florida panthers as population B. The genealogical classes are therefore as follows:

- Model a - pre-rescue Florida panther
- Model b - Backcross with Texas panther
- Model c - F1 hybrid
- Model d - Texas panther
- Model e - Backcross with pre-rescue Florida panther
- Model f - F2 hybrid

Figure 6 shows the posterior probability distribution for post-rescue Florida panthers. Most individuals are a mixture of either genealogical classes **e** and **f** (backcross with pre-rescue Florida panther and F2 hybrid) or **a** and **e** (pre-rescue Florida panther and backcross with pre-rescue Florida panther). Individuals AFP1 and AFP2 (known F1 hybrids) have posterior probabilities for genealogical class **c** (F1 hybrid) of 0.818 and 0.416, respectively, which are too low to warrant assignment. Based on a classification threshold of 0.95 only 7 of 31 individuals were assigned to specific genealogical classes. Table 2 describes the genealogical class assignments for these 7 post-rescue Florida panthers. Four of them are classified as F2 hybrids (Individuals: AFP10, AFP14, AFP24, AFP6), two as backcrosses with pre-rescue Florida panther (Individuals: AFP25, AFP3) and one as pre-rescue Florida panther (Individual: AFP29). For direct comparison, we applied Mongrail to this dataset (see Supplementary Section S3, Figures S14,S15). Results were largely consistent across methods, except for four individuals AFP18 (Figures S14), AFP11 (Figures S14), AFP29 (Figures S15) and AFP5 (Figures S15).

**Table 2.** Assignment of 7 post-rescue Florida panthers.

| Individual Index | Posterior Probability | Genetic Identification | Chosen Model |
| --- | --- | --- | --- |
| AFP10 | 0.995 | F2 | f |
| AFP14 | 0.953 | F2 | f |
| AFP24 | 0.964 | F2 | f |
| AFP25 | 0.992 | Backcross with pre-rescue Florida panther | e |
| AFP29 | 0.999 | pre-rescue Florida panther | a |
| AFP3 | 0.965 | Backcross with pre-rescue Florida panther | e |
| AFP6 | 0.988 | F2 | f |

**Fig. 6.**
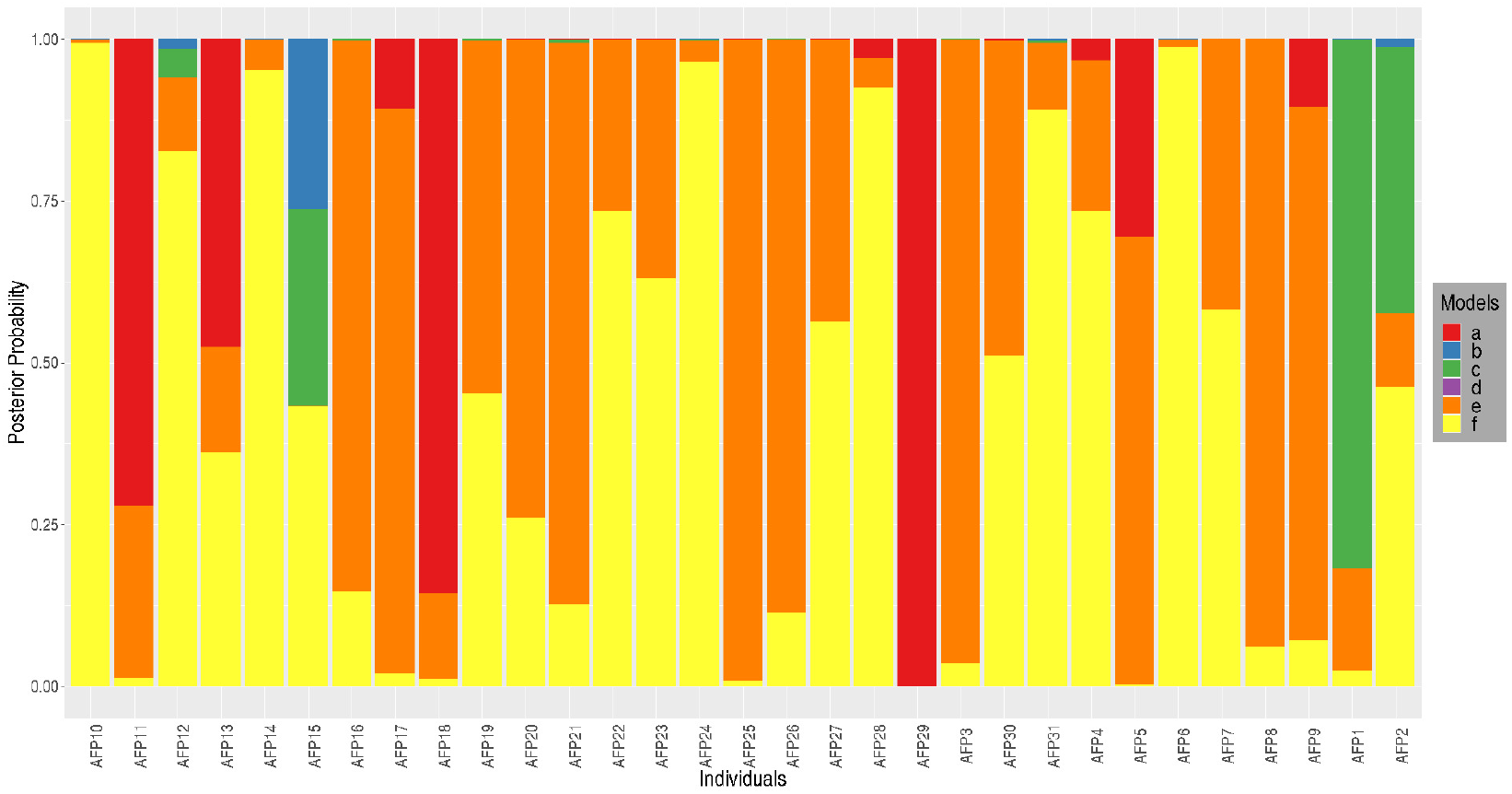
Stacked bar plot showing the distribution of posterior probabilities for 31 post-rescue Florida panthers. The genealogical classes are : **a** - pre-rescue Florida panther, **b** - Backcross with Texas panther, **c** - F1 hybrid, **d** - Texas panther, **e** - Backcross with pre-rescue Florida panther and **f** - F2 hybrid.

## DISCUSSION

In studying the role of hybridization in any context (e.g., conservation genetics, speciation, etc) a necessary first step is to reliably identify hybrids. In our earlier paper (Chakraborty and Rannala, 2023) we developed a Bayesian method (implemented in the program Mongrail) to infer recent hybrids using genomic data. One assumption of Mongrail was that the population haplotype frequencies are known fixed parameters. For empirical datasets population haplotype frequencies are unknown; Chakraborty and Rannala (2023) estimated these using the posterior mean frequency (ignoring uncertainty of haplotype frequencies). Here we address this limitation and develop theory that allows for uncertainty of population haplotype frequencies by integrating over the posterior density using a conjugate prior. We further employed simulation analyses to compare the statistical performance of the two methods. We find that even relatively small samples can produce high posterior probabilities of assignments. The estimates obtained either by using posterior mean point estimates of haplotype frequencies or by integrating over the posterior density of frequencies conditional on the sample are highly similar unless the population sizes are very small, suggesting the previous approximation using estimated haplotype frequencies is fine in many cases. When comparing the new method (implemented in Mongrail 2.0) to the original Mongrail we observed greater uncertainty (represented by more probability associated with alternative genealogical classes) in the posterior probabilities (see Figure 1 and Figure 2). This outcome aligns with our expectation that a method integrating over the posterior density should exhibit more uncertainty than one treating haplotype frequencies as known. Moreover, the uncertainty in Mongrail 2.0 diminishes as the sample size increases, resulting in higher posterior probabilities for the correct genealogical classes. The fact that simulated haplotype counts analyzed using Mongrail 2.0 approach the expected counts under the true population haplotype frequency distribution is indicated by results for *N* = 1000, where the two methods produce essentially similar posterior probabilities.

The performance of both methods is impacted when the number of distinct haplotypes per chromosome (*h*) increases (from 5 to 15) for smaller sample sizes (*N* = 10) (see Figure 3). This is expected, as small sample sizes do not yield reliable estimates of haplotype count data when the number of multinomial categories is large. Additionally, as model complexity increases (purebred → F1 hybrid→ backcross → F2 hybrid) the performance of both methods declines. Qualitatively the new method implemented in Mongrail 2.0 preserves the positive attributes of Mongrail. Specifically, the statistical power of Mongrail 2.0 increases with increasing recombination frequency (map-length of a chromosome) and increasing numbers of chromosomes.

This paper also presents a new algorithm to marginalize over haplotypes in order to calculate the probability of recombinant haplotypes; this is a corrected version of the algorithm described in our earlier paper (Chakraborty and Rannala, 2023). In our earlier approach, we only treated contiguous markers from the same population within a haplotype as jointly distributed, but in fact, all markers inherited from a given parental population should be treated as jointly distributed. This distinction is crucial, as it accurately reflects how meiosis and recombination generate recombinant haplotypes. Multiplying the marginal frequencies of contiguous markers from the same population incorrectly assumes that these segments are independent, contrary to biological reality. Instead contiguous markers derived from the same parental population are inherited together and using the joint frequency of co-transmitted markers correctly captures the ancestry of chromosomal segments formed by recombination. This revised algorithm performs the correct calculation with little additional computational expense. To illustrate this difference, we present a toy example in Supplementary Section S4, comparing the original Mongrail and updated Mongrail 2.0 algorithms. Both methods yield identical results when populations are in linkage equilibrium, which makes sense since the markers are effectively independent in that scenario. The difference between the two algorithms should be insignificant except in the case of large chromosomes that have experienced multiple recombination events. This is observed in Figure S16 (Supplementary Section S5) where Mongrail 2.0 places higher posterior probability on the true genealogical class (model **f**) by comparison with Mongrail for all individuals but one (i7). This holds true even for individuals that were difficult to resolve (i2, i3, i4, i6, i9). Posterior probabilities for individual i7 under genealogical class **f** match between methods to the second decimal place.

We explored Mongrail 2.0’s performance on empirical datasets by applying it to analyze a range of data types comprising non-model organisms: kiwifruit, lizard and puma. Each dataset has unique features that present different challenges for hybrid inferences. The kiwifruit dataset is a genomic study investigating the evolutionary consequences of homoploid hybridization and testing for reduced fitness of hybrids. The lizard dataset includes individuals that were sampled along a linear transect to study the clinal nature of the hybrid zone and the relationship between physical distance and genealogical class. The puma genomic samples were analyzed to study the genomic impact of genetic rescue of Florida panthers through introduction of Texas pumas and exemplify a recent hybridization event with known characteristics and source populations.

In the kiwifruit dataset Mongrail 2.0 classified 7 of 11 presumed *A. zhejiangensis* as F1 hybrids which supports the previous observation that F1 hybrids are more frequent in the hybrid zone (Yu *et al*., 2023). We identified one individual (individual index: 7) as a backcross with *A. eriantha* (genealogical class **b**). A prior Newhybrids analysis identified the same individual (LSZJ-1) as a backcross with *A. eriantha* (Yu *et al*., 2023). By contrast, we identified two individuals (individual index: 1, 5) as F2 hybrids that were previously identified as F1 hybrids (ASX-1 and JNZJ-5) (Yu *et al*., 2023). Our results suggest that hybridization between *Actinidia eriantha* and *Actinidia hemsleyana* may not be a dead-end as previously assumed (Yu *et al*., 2023). The presence of backcross and F2 hybrids contradicts the claim that F1 hybrids are infertile (Yu *et al*., 2023). A possible reason for the lack of F2 individuals in the previous study may be Newhybrids tendency to inflate posterior probabilities of the F1 genealogical class (Chakraborty and Rannala, 2023).

Analysis of the lizard dataset using Mongrail 2.0 classified only 6 of 60 individuals with high posterior probability. Posterior probabilities are typically highly variable, often spread over F2 and backcrosses. There are several factors potentially affecting the method’s ability to infer hybrids confidently. For example, Mongrail 2.0 is based on the assumption that the two populations have been interbreeding for 2 generations. Unclassified hybrids may be an outcome of backcrossing exceeding two generations. The small number of chromosomes analyzed (11 chromosomes) and small sample sizes for the parental populations might also have reduced the power. The power of Mongrail is known to increase with increasing number of chromosomes (Chakraborty and Rannala, 2023).

We also observed (Figure 5) a clear gradient in the proportion of assignment to genealogical classes as we move from north (Fivemile Wash) to south (Snowflake). This sharp transition south of Canoncito is expected as this site is located near the center of the hybrid zone. Mongrail 2.0 captures this spatial pattern which is a characteristic of clinal hybrid zone. The assignment probabilities for individuals from populations north of Canoncito suggests increased Holbrook ancestry as the prominent color is blue (genealogical class **b**: backcross with Holbrook). Whereas individuals from Sevenmile Draw and Snowflake have increased Show Low ancestry as the prominent colors are red (genealogical class **a**: Show Low) and orange (genealogical class **e**: backcross with Show Low). This clear transition in posterior probabilities of assignments is consistent with the findings of Leaché *et al*. (2025).

For the puma dataset, Mongrail 2.0 was able to infer genealogical classes with great certainty for 7 out of 31 post-rescue Florida panthers. Six of these were classified as F2 or backcrosses with pre-rescue Florida panther and one (individual index: 20) was classified as pre-rescue Florida panther. The remaining individuals have little Texas ancestry; the stacked bar plot (Figure 6) rarely exhibits genealogical classes **b** and **d**. These results provide an important insight into the genetic makeup of the post-rescue Florida panthers. They support the findings of Aguilar-Gómez *et al*. (2025) that genetic swamping due to the introduction is unlikely to have occurred. Mongrail 2.0 did not produce high posterior probability for the two known F1 hybrids (individuals AFP1 and AFP2) assigning only 81.8% and 41.3% posterior probability to genealogical class **c** (F1 hybrid). These low posterior probabilities may be due to small sample sizes for the parental populations (5 Texas pumas and 4 pre-rescue Florida panthers).

To allow direct comparison, we applied both Mongrail and Mongrail 2.0 across all empirical datasets (see Supplementary Section S3, Figures S7-S15). Results were largely consistent across methods, except for one individual in the lizard dataset and four individuals in the puma dataset. Observed discrepancies in these cases may be due to small parental population sample sizes.

In summary, we have introduced an improved Bayesian algorithm for inferring hybrids and backcrosses across two generations using sampled genomes from two populations. This new algorithm, Mongrail 2.0, offers two major advancements over the original Mongrail (Chakraborty and Rannala, 2023). First, it relaxes the assumption of known population haplotype frequencies. Mongrail used the Multinomial-Dirichlet posterior mean from reference samples to estimate unknown population haplotype frequencies, disregarding uncertainty. In contrast, Mongrail 2.0 addresses this issue by using the *posterior predictive distribution* which integrates over uncertainties of population allele frequencies. Second, Mongrail 2.0 corrects the probability calculation of recombinant haplotypes by properly marginalizing over haplotypes. These two improvements come with little additional computational cost and simulations show that Mongrail 2.0 performs as effectively as Mongrail in generating high posterior probabilities for the correct genealogical classes when sample sizes are reasonably large, otherwise posterior probabilities are reduced reflecting the additional uncertainty due to uncertain population allele frequencies as expected. Mongrail 2.0 appears to be a statistically conservative method (having low flase positive rates), particularly when the reference population sample size is small, coupled with a high number of distinct haplotypes per chromosome. This is supported by empirical analyses, notably with the lizard and puma datasets, where Mongrail 2.0 faced challenges in inferring hybrids with high certainty. Despite these challenges, Mongrail 2.0 still incorporates linkage and recombination into its model, preserving both power and efficiency even with just 10 markers per chromosome. This speaks to its reliability, especially when the method produces high posterior probabilities. The diverse hybridizing non-model organism datasets used in this study demonstrate the broad applicability and utility of the new method.

## Supporting information

Supplementary Article

## SOFTWARE

The Open Source C program Mongrail 2.0 is available at https://github.com/mongrail/mongrail2.

## DATA AVAILABILITY

Simulated datasets and scripts used for generating the simulations are available at https://github.com/Mongrail-2-0/simulations. The empirical datasets used in this paper can be obtained from the original authors upon request.

## ACKNOWLEDGEMENTS

This work was supported by National Institutes of Health grant GM123306 and National Science Foundation grant DEB-1754254 to BR.

## AUTHOR CONTRIBUTIONS

SC and BR co-developed the theory and co-wrote the manuscript. SC developed the Mongrail 2.0 software and performed the simulation study. SC conducted the analysis of the empirical datasets.

## Appendix A Recombinant haplotype probability

Following Chakraborty and Rannala (2023), an ancestry vector for a haplotype is a boolean array defined as **z** = {*z*_*j*_} where *z*_*j*_ ∈ {0, 1} with 0 or 1 indicating that a SNP locus at position *j* originates from population A or B, respectively. The probability of a recombinant haplotype 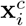 for chromosome *i* is given by

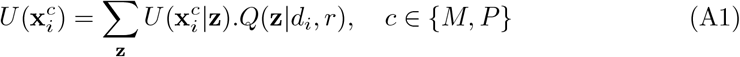

The *Q*(**z** | *d*_*i*_, *r*) term is defined in (Chakraborty and Rannala, 2023) and is restated in Appendix B. The term 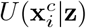 is defined below.

If a haplotype is entirely from either population A or B, the ancestry state **z** is a *L*_*i*_ × 1 vector composed entirely of zeros or ones, respectively. The probability of the haplotype is

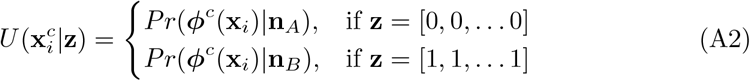

For notational simplicity, we assume *L*_*i*_ = *L*, ∀*i* = 1, 2, … *K* (the argument extends to unequal *L*_*i*_). If the haplotype 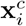 is recombinant between A and B, it can be divided into two sub-haplotypes, 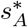 and 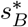 based on ancestry state **z**. A sub-haplotype from population *k* includes only alleles at markers derived from population *k*. Sub-haplotype 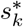 contains alleles of haplotype 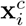 whose markers belong to population *k* ∈ {*A, B*}. The probability of 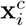 given **z** is a product of sub-haplotype probabilities for 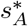 and 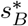. The probability of a sub-haplotype 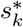 is calculated using the marginal counts of the sub-haplotype obtained by marginalizing over the reference sample haplotype counts (**n**_*k*_) from population *k*. We previously defined matrices of population haplotypes *O*_*A*_ and *O*_*B*_ having dimensions *H* × *L*, where each row is a distinct biallelic haplotype, *H* is the number of distinct haplotypes and *L* is the number of markers. Let *I* be an indexing set, *I* = {1, 2, … *L*}. For a given ancestry state **z**, let *I*_*k*_ ⊂ *I* denote the set of indexes of markers with ancestry from population *k. I*_*A*_ and *I*_*B*_ satisy the conditions,

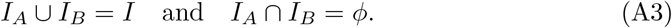

We define a *L* × 1 unit vector 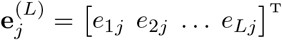 such that for all *j*^*′*^ = 1, 2, …, *L* and *j* ∈ *I*_*k*_

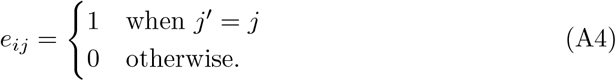

We define the matrix *E*_*k*_ of order *L* × |*I*_*k*_| as

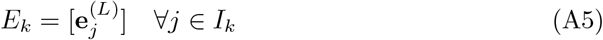

To extract sub-haplotypes whose ancestry belongs to population *k*, we post-multiply *E*_*k*_ with *O*_*k*_

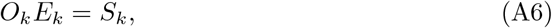

where *S*_*k*_ is of order *H* × |*I*_*k*_|. The rows of the resulting matrix *S*_*k*_ are sub-haplotypes. The set of distinct sub-haplotypes in matrices *S*_*A*_ and *S*_*B*_, denoted by and respectively, 𝒜 and ℬ their corresponding counts (marginals) in the two populations can be found as follows.

Define a *H* × 1 unit vector 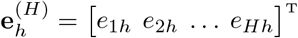 such that for all *h, h*^*′*^ = 1, 2, …, *H*

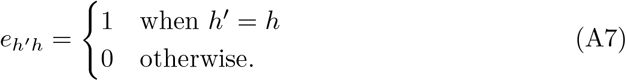

Each of the matrices *S*_*k*_ can be expressed as a column of row vectors

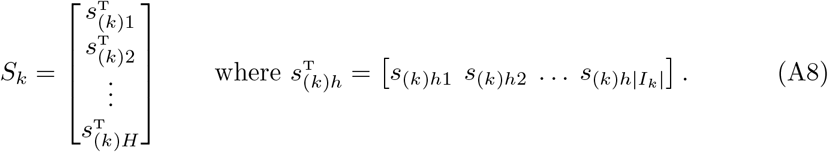

To get the *h*-th row of *S*_*k*_ we perform the operation

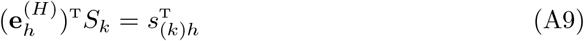

Let *I*_*H*_ be an indexing set, *I*_*H*_ = {1, 2, …, *H*} . For *h*-th row we define a linear transformation,

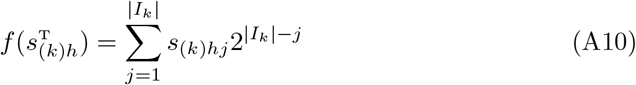

For population A, this linear transformation *f* defines a partition 𝒫_𝒜_ on *S*_*A*_. 𝒫_𝒜_ partitions set *I*_*H*_ into |𝒜| disjoint subsets *A*_1_, *A*_2_, …, *A*_|𝒜|_ where

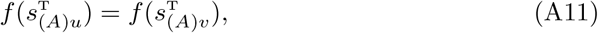

for any *u, v* ∈ *A*_*y*_ satisfying the following three conditions:

1. *A*_*y*_ ≠ *ϕ* for all *y* = 1, 2, …, |𝒜|
2. 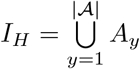
3. *A*_*y*_ ∩ *A*_*y*_^*′*^ = *ϕ* ∀*y* ≠ *y*^*′*^ where *y, y*^′^ = 1, 2, …, |𝒜|

Now we update the corresponding counts of the distinct sub-haplotypes. Let us define the indexing set *Ĩ* = {1, 2, …, |𝒜|}. We can describe a set by associating its element with members of an index set. We define a set *N*_*A*_ that contains the reference sample haplotype counts from population *A*.

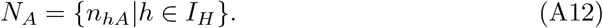

Now bas ed on partition 𝒫_𝒜_ applied on *S*_*A*_, we define set 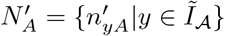 where 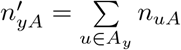.

Therefore we have |𝒜| distinct sub-haplotypes (denoted by 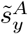 where *y* = 1, 2, … |𝒜| and their corresponding counts denoted by 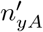). Similarly, we can define a partition 𝒫_*B*_ on *S*_*B*_ which partitions set *I*_*H*_ into |ℬ| disjoint subsets. Subsequently, we have |ℬ| distinct sub-haplotypes (denoted by 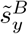 where *y* = 1, 2, … |ℬ| and their corresponding counts denoted by 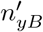).

For *k* ∈ {*A, B*}, the sub-haplotype 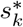 is obtained by post multiplying *E*_*k*_ by 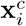

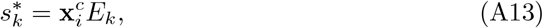

where 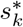 is of order 1 × |*I*_*k*_|. The probability of 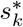 is

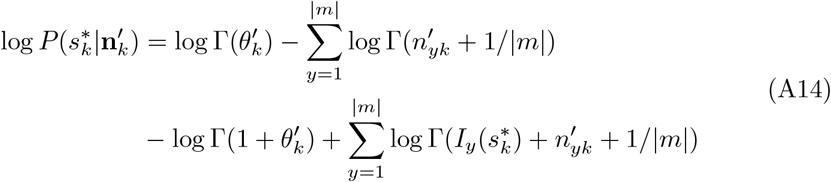

where,

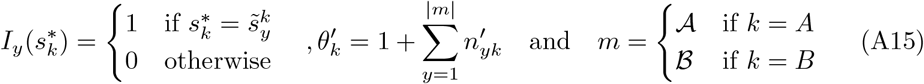

This is equivalent to calculating the posterior predictive distribution of the observed count, 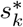, conditioned on 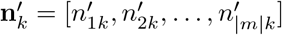. The probability of 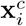 given **z** is

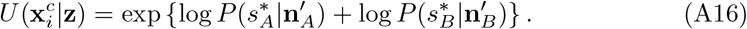

As an example let us consider a case with *L* = 5 markers and *H* = 7 distinct haplotypes. The 7 haplotypes are expressed as rows in the following 7 × 5 matrices:

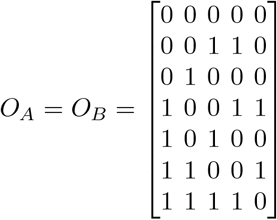

Let the recombinant haplotype be 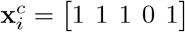 and the ancestry state be **z** = [0 0 1 0 1]. The ancestry state **z**, indicates that markers 1, 2 and 4 belong to population A and markers 3 and 5 belong to population B. The indexing sets are

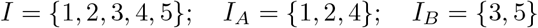

Thus,

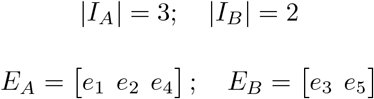

*E*_*A*_ is of order 5 × 3 and *E*_*B*_ is of order 5 × 2. The matrices of the sub-haplotypes are calculated as

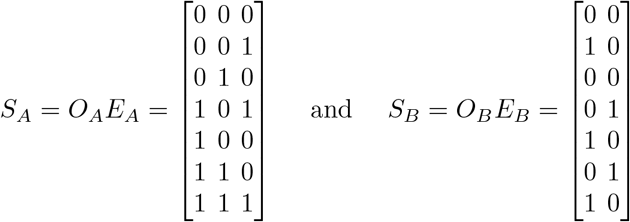

All rows of *S*_*A*_ and 3 rows of *S*_*B*_ are distinct, so the sets of distinct sub-haplotypes are

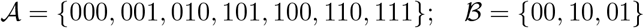

The indexing set is

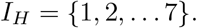

Applying the transformation *f* on *S*_*A*_, partitions the set *I*_*H*_ into |𝒜| = 7 disjoint subsets *A*_*y*_ = {*y*} for all *y* = 1, 2, … 7. Similarly, applying *f* on *S*_*B*_ partitions the set *I*_*H*_ into |ℬ| = 3 disjoint subsets

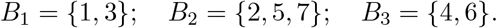

For example, *B*_3_ contains indexes of the rows of *S*_*B*_ (4 and 6) that are identical since

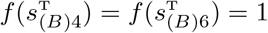

For the set of distinct sub-haplotypes 𝒜 the marginal counts are

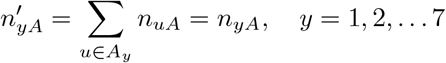

For ℬ, the marginal counts are 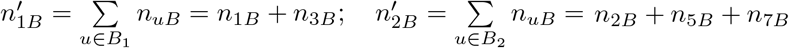 and 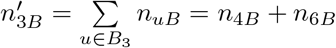. The distinct sub-haplotypes for population A and B are denoted by 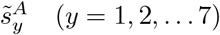 and 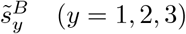. Given the recombinant haplotype 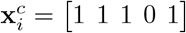 and the ancestry state **z** = [0 0 1 0 1], the sub-haplotypes 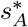 and 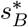 are given by

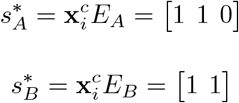

We see that 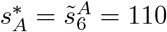 implies 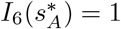 and 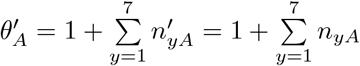.

The log-probability of sub-haplotype 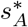 is given by,

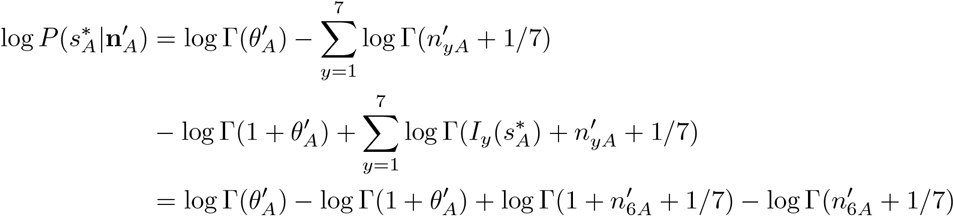

But 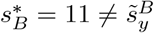 (for any *y* = 1, 2, 3) implies 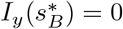 for all *y* = 1, 2, 3 and 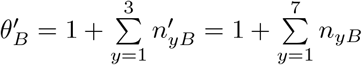. The log-probability of sub-haplotype 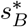 is given by

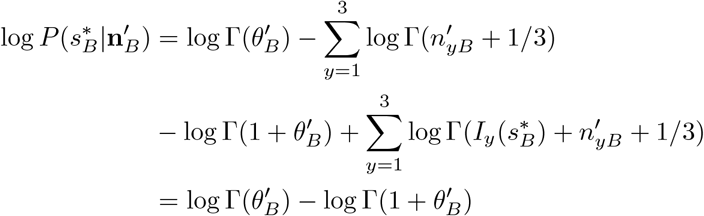

Therefore,

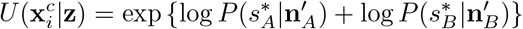

## Appendix B

**Derivation of *Q*(z**|***d***_***i***_, ***r*)**

Before redefining *Q*(**z** | *d*_*i*_, *r*), we first need to define several other terms that will be used in its definition. We already defined in the Theory section (Data and Parameters) that chromosome *i* contains *L*_*i*_ loci with phased biallelic single-nucleotide polymorphisms. Earlier in Appendix A we have also defined an ancestry vector for a haplotype, **z** = {*z*_*j*_} where *z*_*j*_ ∈ {0, 1} denotes the population ancestry state of the marker *j* (where 0 represents population A and 1 indicates the position originates from population B). For chromosome *i*, the physical distance between markers is defined by *d*_*ij*_, where *d*_*ij*_ represents the distance between markers *j* − 1 to *j* and *d*_*i*1_ is the distance from the 5’ end of chromosome *i* to marker 1. We assume a uniform recombination rate *r* on chromosome *i*, measured in centiMorgans (cM) per unit of physical distance. Therefore the map distance between markers *j* − 1 and *j* (for chromosome *i*) is given by *d*_*ij*_ × *r* (cM). Under the assumption of no interference (i.e., recombination events on different intervals are independent of each other) and a uniform recombination rate, recombinations can be modeled as a Poisson process along the chromosome. Thus, the number of recombinations in an interval of length *d*_*ij*_ follows a Poisson distribution with a mean of *rd*_*ij*_. Hence the probability of an even number of recombinations occuring in an interval of length *d*_*ij*_ is given by

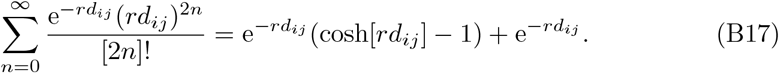

Note that zero recombination (or no recombination) falls under the category of an even number of recombination events. An even number of recombinations results in no change to the ancestry state of the markers. In contrast, an odd number of recombinations between two markers changes the ancestry state of the marker to the right of an interval. Thus, the probability of ancestry change in the interval *d*_*ij*_ is given by

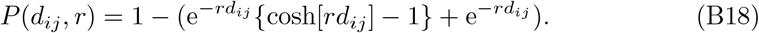

Following the above result, the probability of a particular ancestry state **z** for *L*_*i*_ SNP loci is given by,

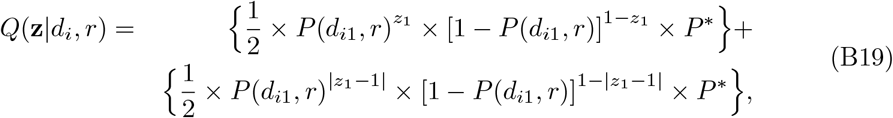

where,

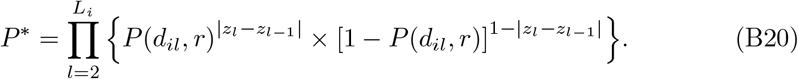

We now explain the derivation of *Q*(**z** | *d*_*i*_, *r*) for a particular ancestry state **z**. The summation in equation B19 of the two terms enclosed in curly braces represents two mutually exclusive and exhaustive events: a chromosome can either be sampled from population A or from population B, each with probability 1/2. Assuming no interference as we move along the chromosome from left to right, transitions from one population ancestry state to another will be calculated as independent conditional probabilities. Consider the first term, where the chromosome is sampled from population A. If *z*_1_ = 0, this indicates that the ancestry state did not change on interval *d*_*i*1_, and the probability of no change is [1 − *P* (*d*_*i*1_, *r*)]. Alternatively, if *z*_1_ = 1, the ancestry state changes on interval *d*_*i*1_, which has a probability of *P* (*d*_*i*1_, *r*). The second term, where the chromosome is sampled from population B, is derived similarly. For the remaining loci (*z*_*l*_; *l* > 1), probabilities are combined into the term *P* ^∗^ defined in equation B20, where *P* (*d*_*il*_, *r*) represents the probability of an ancestry state change (*z*_*l*_ ≠ *z*_*l*−1_), and [1 − *P* (*d*_*il*_, *r*)] represents the probability of no change (*z*_*l*_ = *z*_*l*−1_; *l* ≠ 1).

