## Supplementary Article for "Improved Bayesian inference of hybrids using genome sequences"

### S1 Simulation Analysis

#### S1.1 Simulation Parameters

We reused some of the simulation combinations generated for the Comprehensive Simulation study design in our previous paper Chakraborty and Rannala (2023). To reiterate the simulation design, the parameters (factors) were as follows:

1. Number of chromosomes ($K$)
2. Number of loci per chromosome ($L$)
3. Expected recombination frequency (in cM): $R$ between the first and the last locus
4. Number of distinct haplotype sequences per chromosome for each population ($h$)
5. Allelic configurations of haplotypes, generated by simulating the switches between allele states (see description below). Switch rates used ($c$)
6. Haplotype frequencies, following a Dirichlet distribution (symmetrical with parameter $\alpha$) (see description below)

We simulate haplotype configurations using a 'switching process' that flips the adjacent state of the marker, mimicking the recombination process along the chromosome. We define $p$ as the probability of a switch from 0 to 1 (or 1 to 0), thus the switch rate on a particular interval is $p = c/L$. Haplotype frequencies are simulated using a symmetrical Dirichlet distribution with parameter $\alpha$. For all simulations, the length of each chromosome was fixed at 240 Mb and the recombination rate was set to 1.2 cM/Mb respectively.

#### S1.2 ROC curve analysis

In this paper we considered simulated datasets under the following set of simulation parameters $K = 20, L = 10, c = 0.1, \alpha = 1$. Our goal is to study the performance of the two methods (Mongrail and Mongrail 2.0) under different values of $R$ (1cM or 50cM) and $h$ (5 or 15) for different values of Multinomial sample counts $N = 10, 100, 100$. For each of these 12 simulation combinations (2 for $R$, 2 for $h$, 3 for $N$) we plot the ROC (Receiver Operating Characteristics) curve to examine the methods' power to detect genealogical classes relative to Type I error at different classification thresholds (based on posterior probabilities). The true positive rate (equivalent to power, or sensitivity) is plotted against the false positive rate (equivalent to Type I error, or 1-specificity) to create the ROC curve for the $g$-th genealogical class. The proportion of individuals simulated under the $g$-th genealogical class and classified as such defines the true positive rate. The proportion of individuals simulated under another class but incorrectly classified as the $g$-th class defines the false positive rate. For each of the six genealogical classes, we overlay the ROC curves of Mongrail and Mongrail 2.0 (Figures S1-S6) to facilitate comparisons between the two methods. It should be noted that for Mongrail, posterior probabilities were computed using the posterior mean from the sample counts.

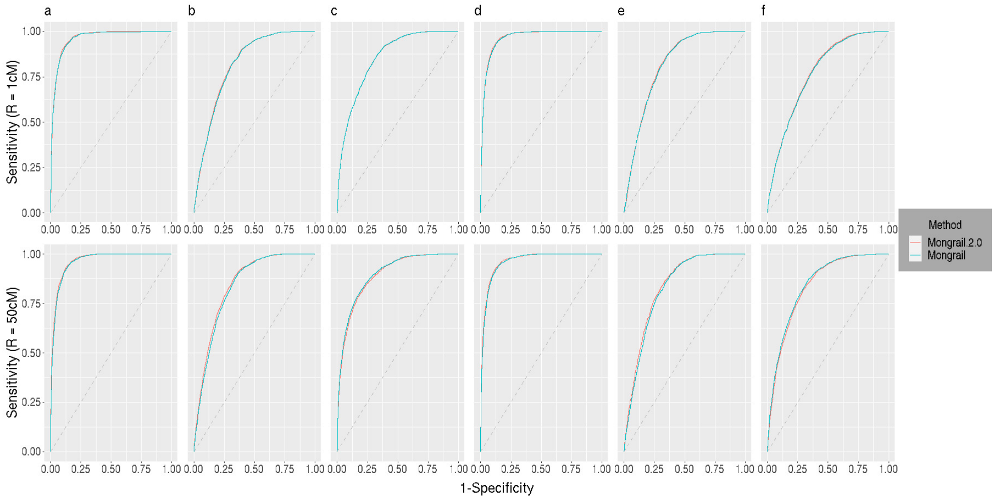


**Fig. S1** Reciever Operator Characteristics (ROC) curves for Mongrail 2.0 (red line) and Mongrail (blue line). The plot is based on 10,000 individuals simulated using parameters: $K = 20, L = 10, h = 5, c = 0.1, \alpha = 1$ with either an expected recombination frequency of $R =$ 1cM (top row) or $R =$ 50cM (bottom row). For Mongrail 2.0, a multinomial sample count of $N = 10$ was used to generate haplotypes for the reference populations. Results for the six genealogical classes (**a**-**f**) are shown from left to right in both rows. The 6 genealogical classes are as follows: **a**-pure population B, **b**-backcross with population A, **c**-F1 hybrid, **d**-pure population A, **e**-backcross with population B, **f**-F2 hybrid.

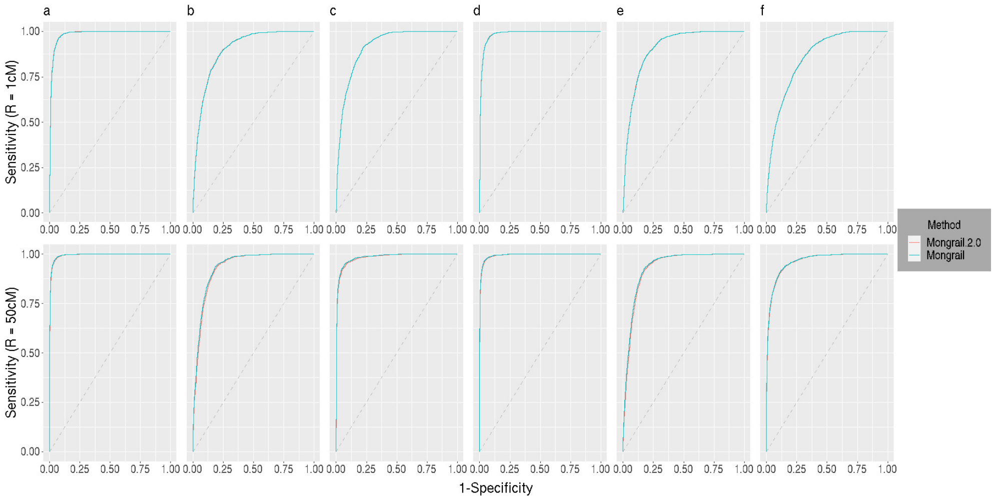


**Fig. S2** Reciever Operator Characteristics (ROC) curves for Mongrail 2.0 (red line) and Mongrail (blue line). The plot is based on 10,000 individuals simulated using parameters: $K = 20, L = 10, h = 5, c = 0.1, \alpha = 1$ with either an expected recombination frequency of $R = 1$cM (top row) or $R = 50$cM (bottom row). For Mongrail 2.0, a multinomial sample count of $N = 100$ was used to generate haplotypes for the reference populations. Results for the six genealogical classes (**a**-**f**) are shown from left to right in both rows. The 6 genealogical classes are as follows: **a**-pure population B, **b**-backcross with population A, **c**-F1 hybrid, **d**-pure population A, **e**-backcross with population B, **f**-F2 hybrid.

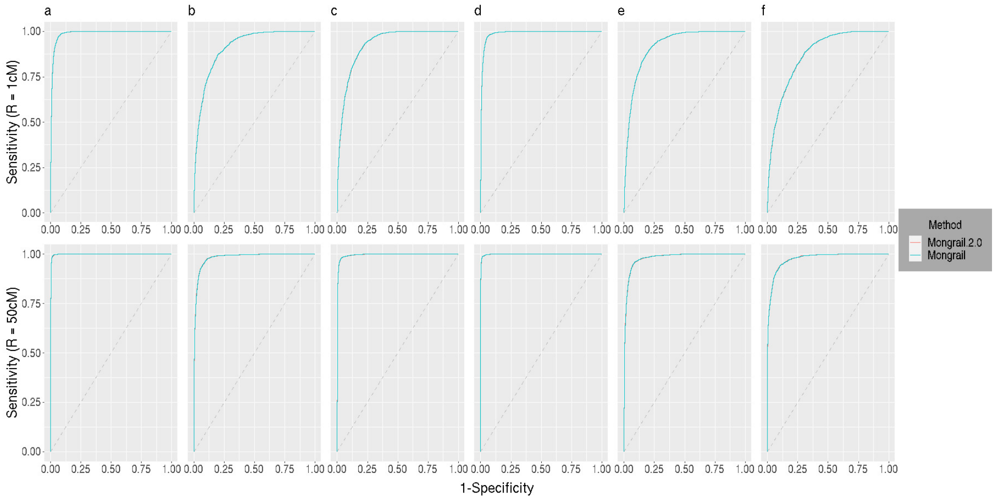


**Fig. S3** Reciever Operator Characteristics (ROC) curves for Mongrail 2.0 (red line) and Mongrail (blue line). The plot is based on 10,000 individuals simulated using parameters: $K = 20, L = 10, h = 5, c = 0.1, \alpha = 1$ with either an expected recombination frequency of $R = 1$cM (top row) or $R = 50$cM (bottom row). For Mongrail 2.0, a multinomial sample count of $N = 1000$ was used to generate haplotypes for the reference populations. Results for the six genealogical classes (**a**-**f**) are shown from left to right in both rows. The 6 genealogical classes are as follows: **a**-pure population B, **b**-backcross with population A, **c**-F1 hybrid, **d**-pure population A, **e**-backcross with population B, **f**-F2 hybrid.

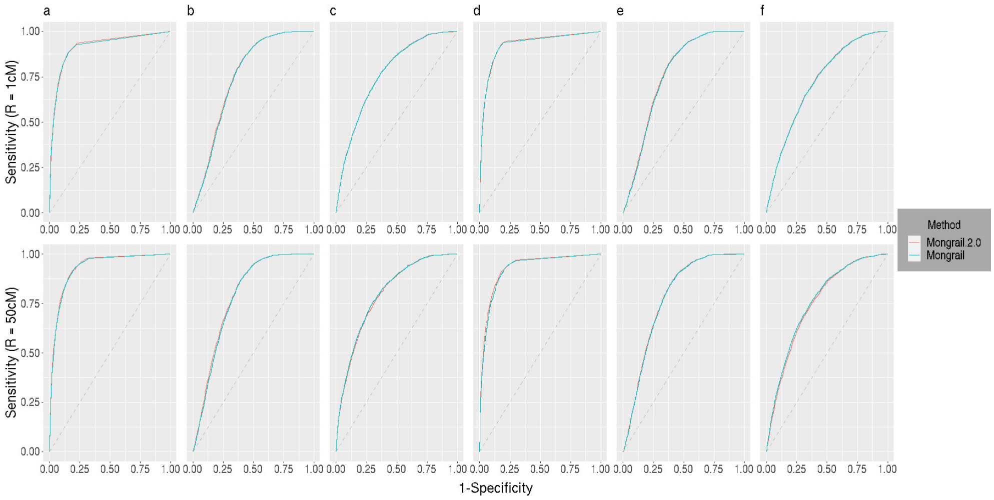


**Fig. S4** Reciever Operator Characteristics (ROC) curves for Mongrail 2.0 (red line) and Mongrail (blue line). The plot is based on 10,000 individuals simulated using parameters: $K = 20, L = 10, h = 15, c = 0.1, \alpha = 1$ with either an expected recombination frequency of $R$ = 1cM (top row) or $R$ = 50cM (bottom row). For Mongrail 2.0, a multinomial sample count of $N$ = 10 was used to generate haplotypes for the reference populations. Results for the six genealogical classes (**a**-**f**) are shown from left to right in both rows. The 6 genealogical classes are as follows: **a**-pure population B, **b**-backcross with population A, **c**-F1 hybrid, **d**-pure population A, **e**-backcross with population B, **f**-F2 hybrid.

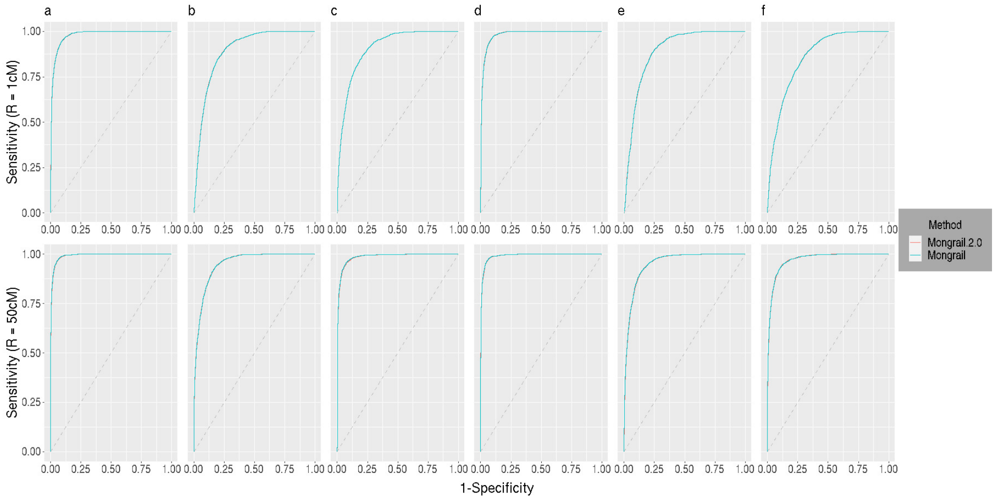


**Fig. S5** Reciever Operator Characteristics (ROC) curves for Mongrail 2.0 (red line) and Mongrail (blue line). The plot is based on 10,000 individuals simulated using parameters: $K = 20, L = 10, h = 15, c = 0.1, \alpha = 1$ with either an expected recombination frequency of $R = 1$cM (top row) or $R = 50$cM (bottom row). For Mongrail 2.0, a multinomial sample count of $N = 100$ was used to generate haplotypes for the reference populations. Results for the six genealogical classes (**a**-**f**) are shown from left to right in both rows. The 6 genealogical classes are as follows: **a**-pure population B, **b**-backcross with population A, **c**-F1 hybrid, **d**-pure population A, **e**-backcross with population B, **f**-F2 hybrid.

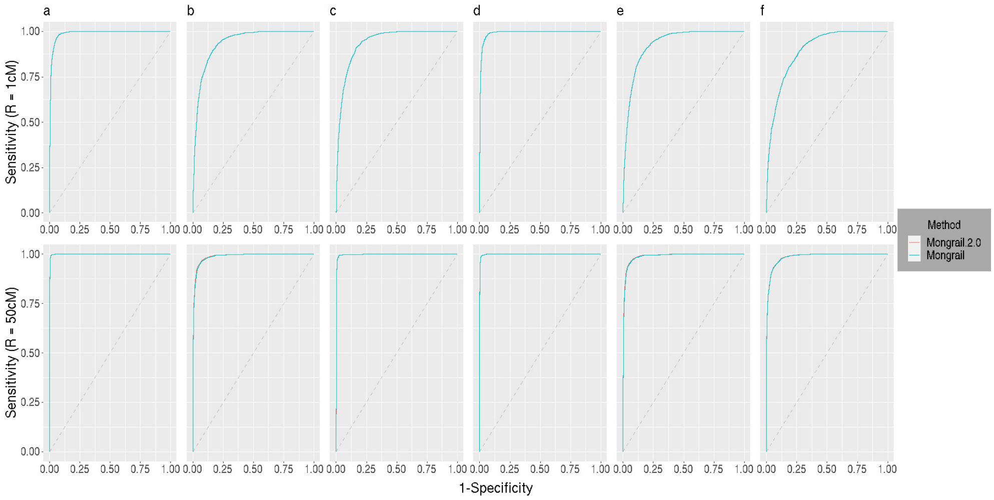


**Fig. S6** Reciever Operator Characteristics (ROC) curves for Mongrail 2.0 (red line) and Mongrail (blue line). The plot is based on 10,000 individuals simulated using parameters: $K = 20, L = 10, h = 15, c = 0.1, \alpha = 1$ with either an expected recombination frequency of $R =$ 1cM (top row) or $R =$ 50cM (bottom row). For Mongrail 2.0, a multinomial sample count of $N = 1000$ was used to generate haplotypes for the reference populations. Results for the six genealogical classes (**a**-**f**) are shown from left to right in both rows. The 6 genealogical classes are as follows: **a**-pure population B, **b**-backcross with population A, **c**-F1 hybrid, **d**-pure population A, **e**-backcross with population B, **f**-F2 hybrid.

### S2 Empirical Analysis

#### S2.1 Kiwifruit

##### S2.1.1 Recombination rate

Recombination rate for either *Actinidia eriantha* or *Actinidia hemsleyana*) were unknown. We extrapolated this value from the linkage map available for another closely related species *Actinidia chinensis*. The genome map length in the female *A. chinensis* was estimated to be 2562 cM (Fraser *et al.*, 2009). The total genome size for all the 29 chromosomes in our dataset was approximately 630 Mb. Thus we used 4.067 cM/Mb ($\frac{2562}{630}$) as the recombination rate for analyzing the kiwifruit dataset.

##### S2.1.2 Recombination frequency and Markers

For each chromosome, we chose recombination frequency (or, map length) such that it's equivalent physical length is approximately equal to the length of the chromosome. This is the "maximally informative" case which was introduced in Chakraborty and Rannala (2023). Then we selected 10 markers from each chromosome by adopting the sliding window approach and then choosing the middlemost window as described in Supplementary Material (Chakraborty and Rannala, 2023).

#### S2.2 Lizard

##### S2.2.1 Recombination rate

Mongrail 2.0 requires recombination rate to be known but this information was not available for plateau fence lizards (*Sceloporus tristichus*). Chakraborty and Rannala (2023) showed that one can use a recombination rate from another closely related species since inference is not sensitive to varying rates. For plateau fence lizards we extrapolated this rate from Eurasian common lizard (*Zootoca vivipara*). Yurchenko *et al.* (2020) calculated the average resolution of linkage maps for *Z. vivipara* males and females to be 0.67 and 0.59 Mb per cM respectively. Since our method requires the recombination rate in units of cM per Mb, we took the inverse of the aforementioned linkage maps and then took an average of the two values. Thus we used a recombination rate of 1.594 cM/Mb ($\frac{0.67^{-1}+0.59^{-1}}{2}$) for our species of study.

##### S2.2.2 Recombination frequency and Markers

Similar to the kiwifruit analysis we considered the "maximally informative" case to choose recombination frequency for each chromosome. We chose 10 markers from each chromosome except chromosome 10. We had only 7 markers available for chromosome 10.

#### S2.3 Puma

##### S2.3.1 Recombination rate

Recombination rate for pumas (*Puma concolor*) was unavailable. But we know that pumas belong to the genus *Felidae* and closely related to domestic cats. Therefore we use 1.9 cM per Mb as the recombination rate for pumas which is the genome wide sex-averaged recombination rate for the autosomes of domestic cats (Li *et al.*, 2016).

##### S2.3.2 Recombination frequency and Markers

Similar to the previous two empirical analyses we considered the "maximally informative" case to choose recombination frequency for each contig. And chose 10 markers from each contig by using the middlemost window approach as mentioned earlier.

### S3 Empirical Analysis: Mongrail versus Mongrail 2.0

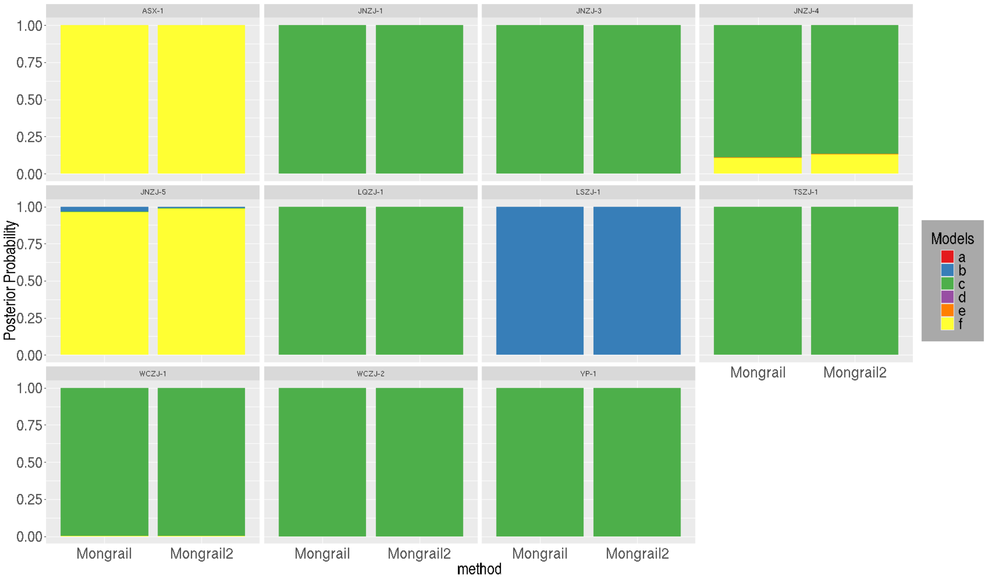


**Fig. S7** Stacked bar plot showing the distribution of posterior probabilities for 11 presumed *A. zhejiangensis* individuals under Mongrail and Mongrail 2.0. The genealogical classes are : **a** - *A. hemsleyana*, **b** - Backcross with *A. eriantha*, **c** - F1 hybrid, **d** - *A. eriantha*, **e** - Backcross with *A. hemsleyana* and **f** - F2 hybrid.

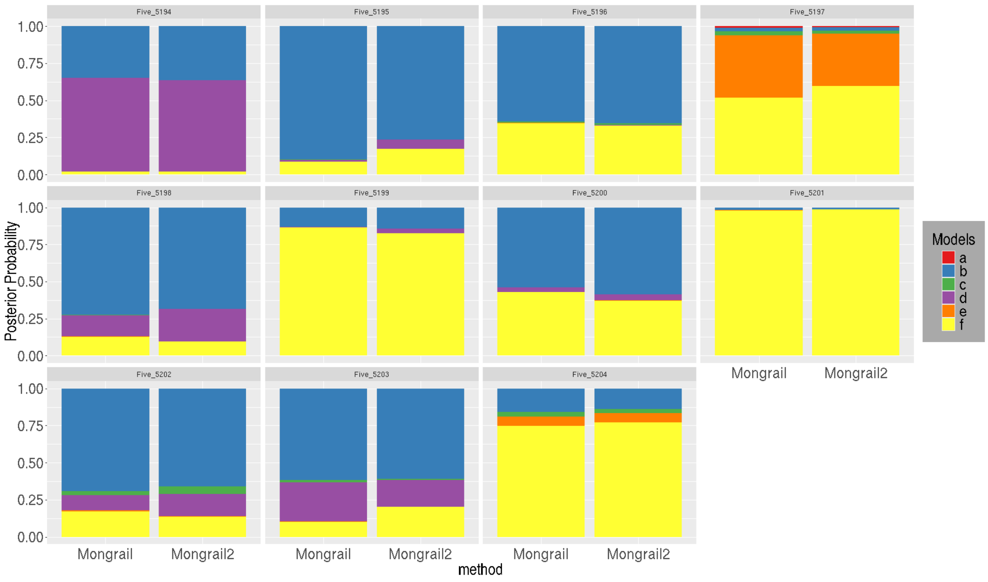


**Fig. S8** Stacked bar plot showing the distribution of posterior probabilities for 11 plateau fence lizards (*Sceloporus tristichus*) sampled from Fivemile Wash under Mongrail and Mongrail 2.0. The genealogical classes are : **a**-pure Show Low, **b**-backcross with Holbrook, **c**-F1 hybrid, **d**-pure Holbrook, **e**-backcross with Show Low, **f**-F2 hybrid.

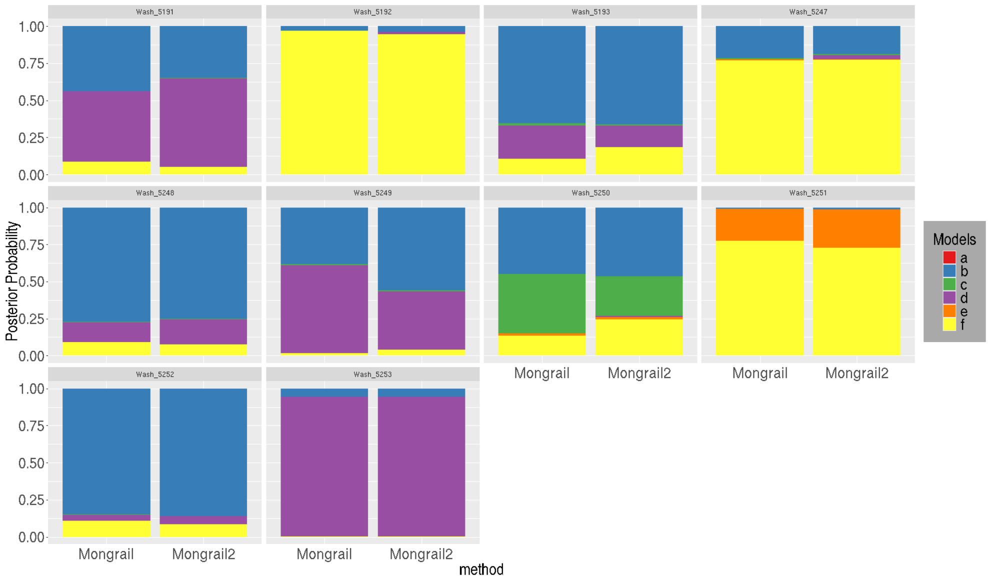


**Fig. S9** Stacked bar plot showing the distribution of posterior probabilities for 10 plateau fence lizards (*Sceloporus tristichus*) sampled from Washboard Wash under Mongrail and Mongrail 2.0. The genealogical classes are : **a**-pure Show Low, **b**-backcross with Holbrook, **c**-F1 hybrid, **d**-pure Holbrook, **e**-backcross with Show Low, **f**-F2 hybrid.

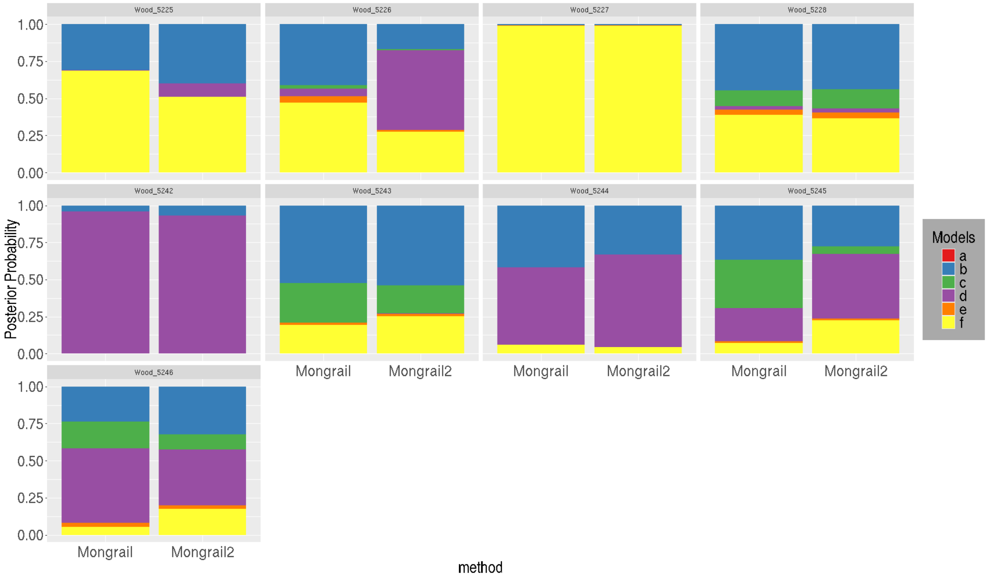


**Fig. S10** Stacked bar plot showing the distribution of posterior probabilities for 9 plateau fence lizards (*Sceloporus tristichus*) sampled from Woodruff under Mongrail and Mongrail 2.0. The genealogical classes are : **a**-pure Show Low, **b**-backcross with Holbrook, **c**-F1 hybrid, **d**-pure Holbrook, **e**-backcross with Show Low, **f**-F2 hybrid.

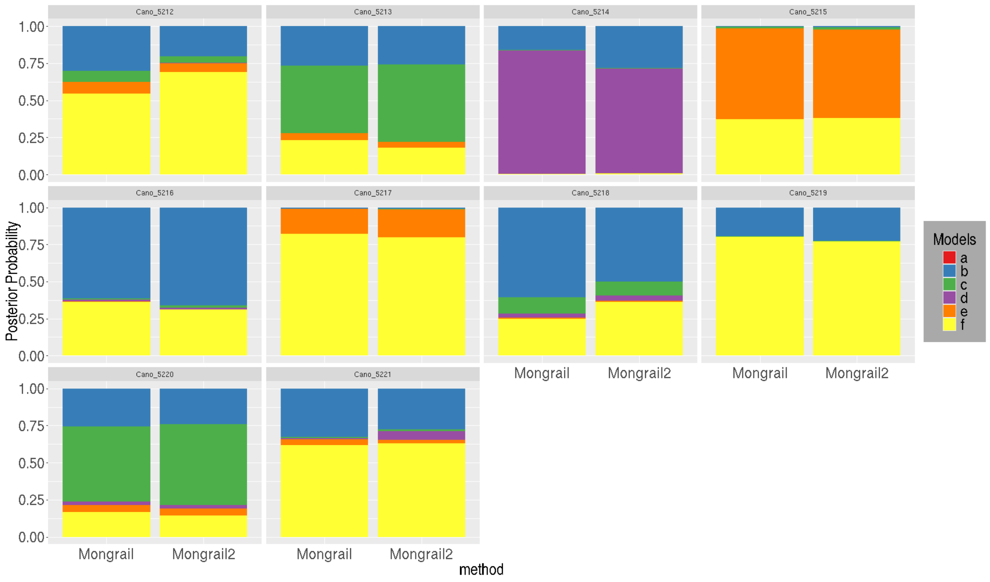


**Fig. S11** Stacked bar plot showing the distribution of posterior probabilities for 10 plateau fence lizards (*Sceloporus tristichus*) sampled from Canoncito under Mongrail and Mongrail 2.0. The genealogical classes are : **a**-pure Show Low, **b**-backcross with Holbrook, **c**-F1 hybrid, **d**-pure Holbrook, **e**-backcross with Show Low, **f**-F2 hybrid.

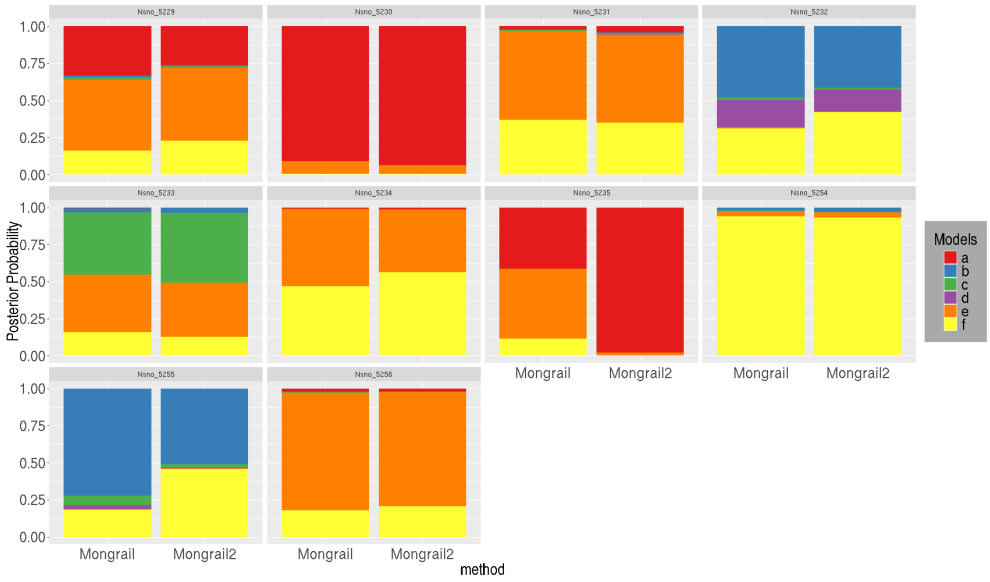


**Fig. S12** Stacked bar plot showing the distribution of posterior probabilities for 10 plateau fence lizards (*Sceloporus tristichus*) sampled from Sevenmile Draw under Mongrail and Mongrail 2.0. The genealogical classes are : **a**-pure Show Low, **b**-backcross with Holbrook, **c**-F1 hybrid, **d**-pure Holbrook, **e**-backcross with Show Low, **f**-F2 hybrid.

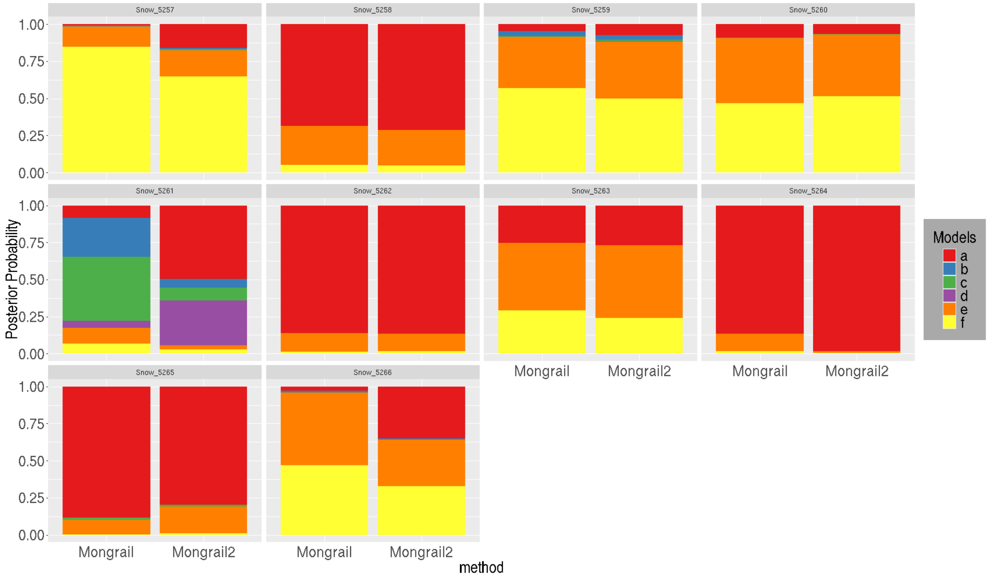


**Fig. S13** Stacked bar plot showing the distribution of posterior probabilities for 10 plateau fence lizards (*Sceloporus tristichus*) sampled from Snowflake under Mongrail and Mongrail 2.0. The genealogical classes are : **a**-pure Show Low, **b**-backcross with Holbrook, **c**-F1 hybrid, **d**-pure Holbrook, **e**-backcross with Show Low, **f**-F2 hybrid.

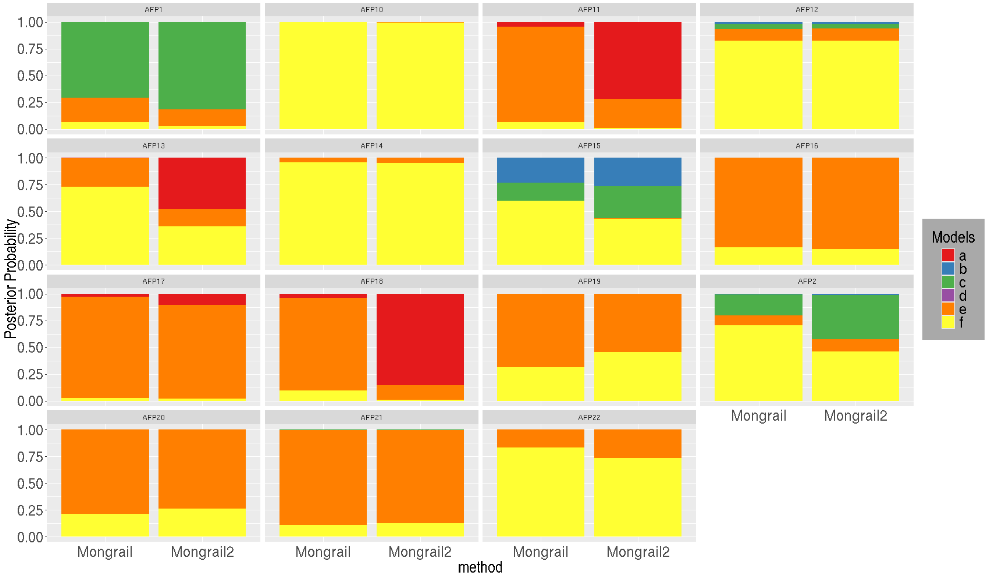


**Fig. S14** Stacked bar plot showing the distribution of posterior probabilities for 15 post-rescue Florida panthers under Mongrail and Mongrail 2.0. The genealogical classes are : **a** - pre-rescue Florida panther, **b** - Backcross with Texas panther, **c** - F1 hybrid, **d** - Texas panther, **e** - Backcross with pre-rescue Florida panther and **f** - F2 hybrid.

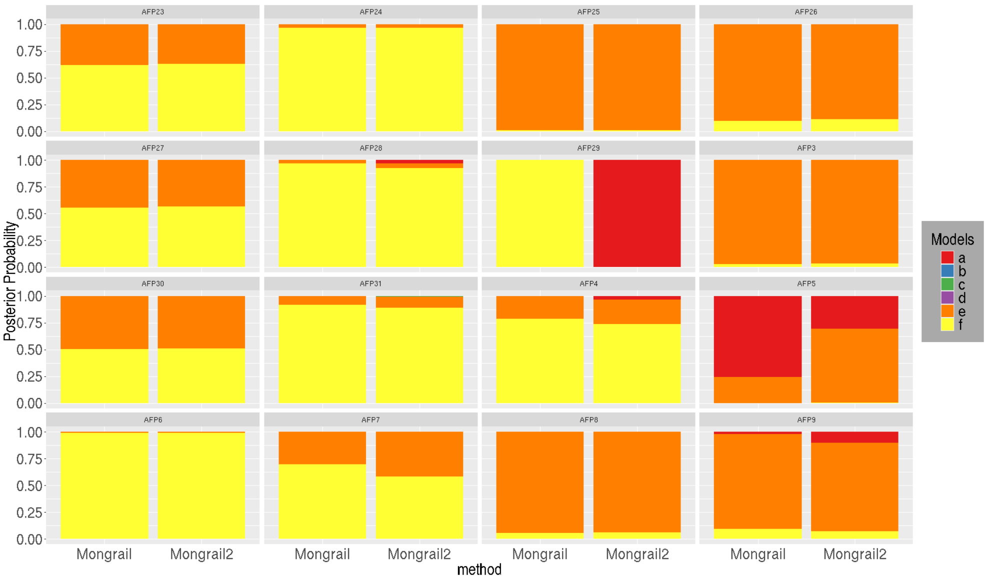


**Fig. S15** Stacked bar plot showing the distribution of posterior probabilities for rest of the 16 post-rescue Florida panthers under Mongrail and Mongrail 2.0. The genealogical classes are : **a** - pre-rescue Florida panther, **b** - Backcross with Texas panther, **c** - F1 hybrid, **d** - Texas panther, **e** - Backcross with pre-rescue Florida panther and **f** - F2 hybrid.

### S4 Correct marginalization of haplotypes

Here, we illustrate how the two methods, Mongrail and Mongrail 2.0, differ in calculating the probability of recombinant haplotypes, using a simple toy example. For clarity, we assume that the population haplotype frequencies for both populations A and B are known. This allows us to focus solely on the differences in the marginalization algorithms used by the two methods, without the added complexity of uncertainty in population haplotype frequencies. We also consider two cases: Linkage Disequilibrium and Linkage Equilibrium within this framework. Let us consider an example with $L = 4$ markers and $H = 16$ distinct haplotypes. The 16 haplotypes are expressed as rows in the following $16 \times 4$ matrices:

$$O_A = O_B = \begin{bmatrix} 0&0&0&0 \\ 0&0&0&1 \\ 0&0&1&0 \\ 0&0&1&1 \\ 0&1&0&0 \\ 0&1&0&1 \\ 0&1&1&0 \\ 0&1&1&1 \\ 1&0&0&0 \\ 1&0&0&1 \\ 1&0&1&0 \\ 1&0&1&1 \\ 1&1&0&0 \\ 1&1&0&1 \\ 1&1&1&0 \\ 1&1&1&1 \end{bmatrix}$$

Let the recombinant haplotype be $\mathbf{x}_i^c = \begin{bmatrix} 1 & 1 & 1 & 0 \end{bmatrix}$ and the ancestry state be $\mathbf{z} = \begin{bmatrix} 0 & 1 & 1 & 0 \end{bmatrix}$. The ancestry state $\mathbf{z}$, indicates that markers 1 and 4 belong to population A and markers 2 and 3 belong to population B.

$$\mathbf{z} = \left[ \underbrace{0}_{A} \quad \underbrace{1 \quad 1}_{B \quad B} \quad \underbrace{0}_{A} \right]$$

In this section we will show how probability of recombinant haplotype $\mathbf{x}_i^c$ given $\mathbf{z}$ is calculated differently in Mongrail $[f(\mathbf{x}_i^c|\mathbf{z})]$ versus Mongrail 2.0 $[U(\mathbf{x}_i^c|\mathbf{z})]$. Let's elucidate how the sub-haplotypes are considered for the two methods.

#### S4.1 Mongrail

Based on $\mathbf{z}$, the recombinant haplotype will be divided into three sub-haplotype segments $\{1\}$, $\{1\ 1\}$ and $\{0\}$ of lengths $j_1 = 1$, $j_2 = 2$ and $j_3 = 1$ respectively.

$$\mathbf{x}_i^c = \left[ \underbrace{1}_{A} \quad \underbrace{1 \quad 1}_{B \quad B} \quad \underbrace{0}_{A} \right]$$

In Mongrail, we obtain the marginal frequency of alleles on these three sub-haplotypes from their corresponding population (color coded) and multiply them to get the final $f(\mathbf{x}_i^c|\mathbf{z})$. Under this setup we defined the marginal frequency of alleles on sub haplotypes $\{x_{iu}^c, x_{i(u+1)}^c, x_{i(u+2)}^c, \ldots, x_{iv}^c\}$ from the $u$-th marker to the $v$-th marker where $1 \leq u < v \leq 4$ in population A (or, B) as $f^A(x_{i[u,v]})$ or, $f^B(x_{i[u,v]})$ respectively. In terms of mathematical notation the probability of the recombinant haplotype $\mathbf{x}_i^c$ given $\mathbf{z}$ is

$$f(\mathbf{x}_i^c|\mathbf{z}) = f^A(x_{i[1,1]}^c) \times f^B(x_{i[2,3]}^c) \times f^A(x_{i[4,4]}^c)$$
$$= f^A(\{1\}) \times f^B(\{11\}) \times f^A(\{0\}) \quad (1)$$

#### S4.2 Mongrail 2.0

Under Mongrail 2.0 any recombinant haplotype $\mathbf{x}_i^c$ is always divided into two sub-haplotypes, $s_A^*$ and $s_B^*$ based on its ancestry state $\mathbf{z}$. Sub-haplotype $s_k^*$ contains alleles of haplotype $\mathbf{x}_i^c$ whose markers belong to population $k \in \{A, B\}$. Therefore for this toy example

$$s_A^* = \begin{bmatrix} 1 & 0 \end{bmatrix}, \qquad s_B^* = \begin{bmatrix} 1 & 1 \end{bmatrix}$$

The blue box around 1st and 4th columns gives us the sub-haplotype matrix $S_A$ for population A and the red box around 2nd and 3rd columns gives the sub-haplotype matrix $S_B$ for population B.

$$O_A = O_B = \begin{bmatrix} 0 & 0 & 0 & 0 \\ 0 & 0 & 0 & 1 \\ 0 & 0 & 1 & 0 \\ 0 & 0 & 1 & 1 \\ 0 & 1 & 0 & 0 \\ 0 & 1 & 0 & 1 \\ 0 & 1 & 1 & 0 \\ 0 & 1 & 1 & 1 \\ 1 & 0 & 0 & 0 \\ 1 & 0 & 0 & 1 \\ 1 & 0 & 1 & 0 \\ 1 & 0 & 1 & 1 \\ 1 & 1 & 0 & 0 \\ 1 & 1 & 0 & 1 \\ 1 & 1 & 1 & 0 \\ 1 & 1 & 1 & 1 \end{bmatrix}, \quad S_A = \underbrace{\begin{bmatrix} 0 & 0 \\ 0 & 1 \\ 0 & 0 \\ 0 & 1 \\ 0 & 0 \\ 0 & 1 \\ 0 & 0 \\ 0 & 1 \\ 1 & 0 \\ 1 & 1 \\ 1 & 0 \\ 1 & 1 \\ 1 & 0 \\ 1 & 1 \\ 1 & 0 \\ 1 & 1 \end{bmatrix}}_{\text{columns 1 \& 4}}, \quad S_B = \underbrace{\begin{bmatrix} 0 & 0 \\ 0 & 0 \\ 0 & 1 \\ 0 & 1 \\ 1 & 0 \\ 1 & 0 \\ 1 & 1 \\ 1 & 1 \\ 0 & 0 \\ 0 & 0 \\ 0 & 1 \\ 0 & 1 \\ 1 & 0 \\ 1 & 0 \\ 1 & 1 \\ 1 & 1 \end{bmatrix}}_{\text{columns 2 \& 3}}$$

The probability of sub-haplotype $s_A^* = \begin{bmatrix} 1 & 0 \end{bmatrix}$ is the marginal frequency of haplotype $\begin{bmatrix} 1 & 0 \end{bmatrix}$ in matrix $S_A$ (summing the frequencies of haplotypes in blue outlined boxes). Similarly, probability of sub-haplotype $s_B^* = \begin{bmatrix} 1 & 1 \end{bmatrix}$ is the marginal frequency of haplotype $\begin{bmatrix}1 & 1\end{bmatrix}$ in matrix $S_B$ (summing the frequencies of haplotypes in red outlined boxes). Therefore probability of the recombinant haplotype $\mathbf{x}_i^c$ given $\mathbf{z}$ is a product of sub-haplotype probabilities for $s_A^*$ and $s_B^*$.

Let us calculate these probabilities for Mongrail and Mongrail 2.0 under two specific cases:

- **Case I: Linkage Disequilibrium**
  Say, population A is in linkage disequilibrium. The arrows (in matrix $O_A$) from each haplotype point to its corresponding frequency ($f_h^A; h = 1, 2, \ldots, 16$). Whereas say population B is at equilibrium so the population frequencies for all 16 haplotypes is 0.0625. $f^B = \{0.0625, 0.0625, \ldots, 0.0625\}$.

$$O_A = \begin{bmatrix} 0 & 0 & 0 & 0 \\ 0 & 0 & 0 & 1 \\ 0 & 0 & 1 & 0 \\ 0 & 0 & 1 & 1 \\ 0 & 1 & 0 & 0 \\ 0 & 1 & 0 & 1 \\ 0 & 1 & 1 & 0 \\ 0 & 1 & 1 & 1 \\ 1 & 0 & 0 & 0 \\ 1 & 0 & 0 & 1 \\ 1 & 0 & 1 & 0 \\ 1 & 0 & 1 & 1 \\ 1 & 1 & 0 & 0 \\ 1 & 1 & 0 & 1 \\ 1 & 1 & 1 & 0 \\ 1 & 1 & 1 & 1 \end{bmatrix} \begin{array}{l} \rightarrow f_1^A = 0.273 \\ \rightarrow f_2^A = 0 \\ \rightarrow f_3^A = 0.019 \\ \rightarrow f_4^A = 0.001 \\ \rightarrow f_5^A = 0 \\ \rightarrow f_6^A = 0.001 \\ \rightarrow f_7^A = 0 \\ \rightarrow f_8^A = 0.104 \\ \rightarrow f_9^A = 0.001 \\ \rightarrow f_{10}^A = 0 \\ \rightarrow f_{11}^A = 0 \\ \rightarrow f_{12}^A = 0.593 \\ \rightarrow f_{13}^A = 0 \\ \rightarrow f_{14}^A = 0.007 \\ \rightarrow f_{15}^A = 0.001 \\ \rightarrow f_{16}^A = 0 \end{array}$$

  Under Mongrail, we follow equation 1 to calculate the probability of recombinant haplotype given ancestry state. The probability of the first sub-haplotype $f^A(\{1\})$ is the marginal frequency of allele 1 (marker 1) in population A which is summing over all the haplotype frequencies that has allele 1 in marker 1 for matrix $O_A$. Thus $f^A(x^c_{i[1,1]}) = \sum_{h=9}^{16} f_h^A = (0.001 + 0 + 0 + 0.593 + 0 + 0.007 + 0.001 + 0) = 0.602$. Similarly, we calculate $f^B(\{11\})$ which is $f^B(x^c_{i[2,3]}) = (f_7^B + f_8^B + f_{15}^B + f_{16}^B) = (0.0625 \times 4) = 0.25$. And frequency of third sub-haplotype $f^A(\{0\})$ is $f^A(x^c_{i[4,4]}) = \sum_{h=0}^{7} f_{2h+1}^A = (0.273 + 0.019 + 0 + 0 + 0.001 + 0 + 0 + 0.001) = 0.294$. Therefore under Mongrail, the probability of recombinant haplotype $\mathbf{x}_i^c$ given $\mathbf{z}$ is

$$f(\mathbf{x}_i^c|\mathbf{z}) = f^A(x^c_{i[1,1]}) \times f^B(x^c_{i[2,3]}) \times f^A(x^c_{i[4,4]})$$

$$
\begin{aligned}
&= f^A(\{1\}) \times f^B(\{11\}) \times f^A(\{0\}) \\
&= (0.602) \times (0.25) \times (0.294) \\
&= 0.044247
\end{aligned} \tag{2}
$$

Under Mongrail 2.0, when we consider matrix $S_A$ ($S_B$) derived from $O_A$ ($O_B$) their corresponding frequencies for each row (sub-haplotype) remain unchanged.

$$
S_A = \begin{bmatrix} 0 & 0 \\ 0 & 1 \\ 0 & 0 \\ 0 & 1 \\ 0 & 0 \\ 0 & 1 \\ 0 & 0 \\ 0 & 1 \\ \boxed{1 \quad 0} \\ 1 & 1 \\ \boxed{1 \quad 0} \\ 1 & 1 \\ \boxed{1 \quad 0} \\ 1 & 1 \\ \boxed{1 \quad 0} \\ 1 & 1 \end{bmatrix}
\begin{array}{l} \rightarrow f_1^A = 0.273 \\ \rightarrow f_2^A = 0 \\ \rightarrow f_3^A = 0.019 \\ \rightarrow f_4^A = 0.001 \\ \rightarrow f_5^A = 0 \\ \rightarrow f_6^A = 0.001 \\ \rightarrow f_7^A = 0 \\ \rightarrow f_8^A = 0.104 \\ \rightarrow f_9^A = 0.001 \\ \rightarrow f_{10}^A = 0 \\ \rightarrow f_{11}^A = 0 \\ \rightarrow f_{12}^A = 0.593 \\ \rightarrow f_{13}^A = 0 \\ \rightarrow f_{14}^A = 0.007 \\ \rightarrow f_{15}^A = 0.001 \\ \rightarrow f_{16}^A = 0 \end{array},
\quad
S_B = \begin{bmatrix} 0 & 0 \\ 0 & 0 \\ 0 & 1 \\ 0 & 1 \\ 1 & 0 \\ 1 & 0 \\ \boxed{1 \quad 1} \\ \boxed{1 \quad 1} \\ 0 & 0 \\ 0 & 0 \\ 0 & 1 \\ 0 & 1 \\ 1 & 0 \\ 1 & 0 \\ \boxed{1 \quad 1} \\ \boxed{1 \quad 1} \end{bmatrix}
\begin{array}{l} \rightarrow f_1^B = 0.0625 \\ \rightarrow f_2^B = 0.0625 \\ \rightarrow f_3^B = 0.0625 \\ \rightarrow f_4^B = 0.0625 \\ \rightarrow f_5^B = 0.0625 \\ \rightarrow f_6^B = 0.0625 \\ \rightarrow f_7^B = 0.0625 \\ \rightarrow f_8^B = 0.0625 \\ \rightarrow f_9^B = 0.0625 \\ \rightarrow f_{10}^B = 0.0625 \\ \rightarrow f_{11}^B = 0.0625 \\ \rightarrow f_{12}^B = 0.0625 \\ \rightarrow f_{13}^B = 0.0625 \\ \rightarrow f_{14}^B = 0.0625 \\ \rightarrow f_{15}^B = 0.0625 \\ \rightarrow f_{16}^B = 0.0625 \end{array}
$$

Thus probability of sub-haplotype $s_A^* = [1\ 0]$ is the marginal frequency of haplotype $[1\ 0]$ in matrix $S_A$. Thus $P(s_A^*) = f_9^A + f_{11}^A + f_{13}^A + f_{15}^A = (0.001 + 0 + 0 + 0.001) = 0.002$. Similarly, probability of sub-haplotype $s_B^* = [1\ 1]$ is the marginal frequency of haplotype $[1\ 1]$ in matrix $S_B$. Thus $P(s_B^*) = f_7^B + f_8^B + f_{15}^B + f_{16}^B = (0.0625 \times 4) = 0.25$ Therefore under Mongrail 2.0, probability of the recombinant haplotype $\mathbf{x}_i^c$ given $\mathbf{z}$ is

$$
\begin{aligned}
U(\mathbf{x}_i^c|\mathbf{z}) &= P(s_A^*) \times P(s_B^*) \\
&= (0.002) \times (0.25) \\
&= 0.0005
\end{aligned} \tag{3}
$$

- **Case II: Linkage Equilibrium** Both the populations are at equilibrium. We consider population frequencies for all 16 haplotypes to be 0.0625. $f^A = f^B = \{0.0625, 0.0625, \ldots, 0.0625\}$. Then all the previous formulae remain the same except the values for population A. Therefore under Mongrail, the probability of recombinant haplotype $\mathbf{x}_i^c$ given $\mathbf{z}$ is

$$
\begin{aligned}
f(\mathbf{x}_i^c|\mathbf{z}) &= f^A(x_{i[1,1]}^c) \times f^B(x_{i[2,3]}^c) \times f^A(x_{i[4,4]}^c) \\
&= f^A(\{1\}) \times f^B(\{11\}) \times f^A(\{0\}) \\
&= (0.0625 \times 8) \times (0.0625 \times 4) \times (0.0625 \times 8) \\
&= 0.0625
\end{aligned} \tag{4}
$$

Therefore under Mongrail 2.0, probability of the recombinant haplotype $\mathbf{x}_i^c$ given $\mathbf{z}$ is

$$
\begin{aligned}
U(\mathbf{x}_i^c|\mathbf{z}) &= P(s_A^*) \times P(s_B^*) \\
&= (0.0625 * 4) \times (0.0625 * 4) \\
&= 0.0625
\end{aligned} \tag{5}
$$

Thus we see that Mongrail and Mongrail 2.0 produces the same results if both populations are at linkage equilibrium. But the two methods differ if at least one of the populations is in linkage disequilibrium.

### S5 Mongrail 2.0 versus Mongrail

We construct a simple example with $L = 10$ markers and $H = 2$ distinct haplotypes for both populations A and B. The two haplotypes are expressed as rows in the following $2 \times 10$ matrices $O_A$ and $O_B$. The arrows (in matrix $O_A$ and $O_B$) from each haplotype point to its corresponding true frequency ($f_h^A$ and $f_h^B; h = 1, 2$).

$$
O_A = \begin{bmatrix} 0 & 0 & 0 & 0 & 0 & 0 & 0 & 0 & 0 & 0 \\ 1 & 1 & 1 & 1 & 1 & 1 & 1 & 1 & 1 & 1 \end{bmatrix} \begin{matrix} \longrightarrow f_1^A = 0.9 \\ \longrightarrow f_2^A = 0.1 \end{matrix}
$$

$$
O_B = \begin{bmatrix} 0 & 0 & 0 & 0 & 0 & 0 & 0 & 0 & 0 & 0 \\ 1 & 1 & 1 & 1 & 1 & 1 & 1 & 1 & 1 & 1 \end{bmatrix} \begin{matrix} \longrightarrow f_1^B = 0.1 \\ \longrightarrow f_2^B = 0.9 \end{matrix}
$$

We consider $K = 4$ chromosomes for both populations with the same haplotypes and frequencies. For each chromosome we choose a size of 240 Mb and a recombination rate of 1.2 cM/Mb. We choose $L = 10$ markers on the chromosome in a way such that the distance between the first and last marker is 282 cM. This choice allows for more recombination events to take place per meiosis. We simulate 10 individuals from genealogical class $\mathbf{f}$ (F2 hybrid) based on the true haplotype frequencies.

Now based on these true haplotype frequencies we consider a multinomial sample size of $N = 50$ to generate reference population datasets. We applied Mongrail 2.0 to these 10 simulated individuals using the simulated reference samples to compute the posterior probabilities. We also applied Mongrail to the same individuals to compute posterior probabilities using the posterior mean from the sample counts. The results of the analysis are presented in Figure S16.

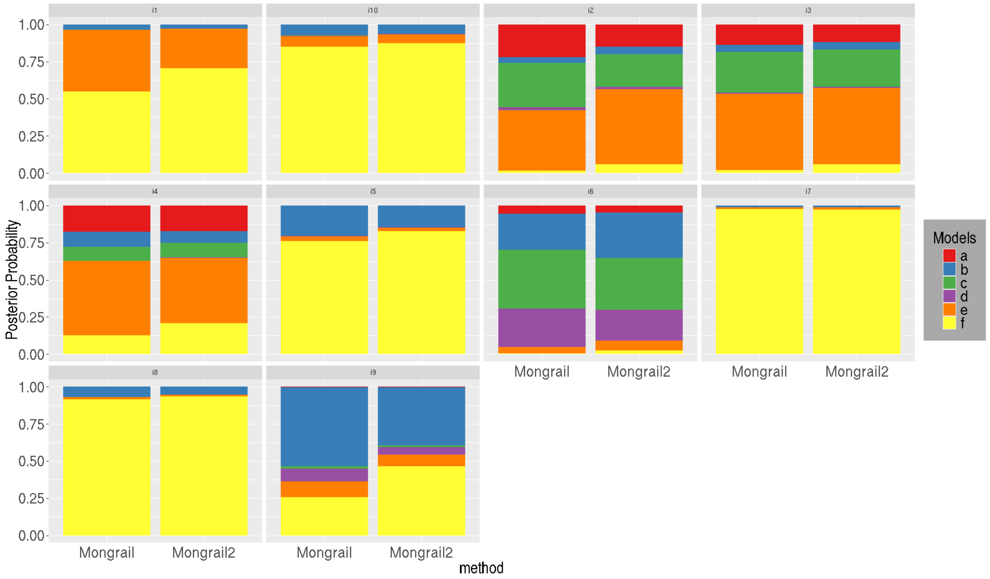


**Fig. S16** Stacked bar plot showing the distribution of posterior probabilities for 10 simulated individuals (generated under genealogical class **f**) under Mongrail and Mongrail 2.0. The genealogical classes are **a**-pure population B, **b**-backcross with population A, **c**-F1 hybrid, **d**-pure population A, **e**-backcross with population B, **f**-F2 hybrid.
